# Inter-generational nuclear crosstalk links the control of gene expression to programmed genome rearrangements during the *Paramecium* sexual cycle

**DOI:** 10.1101/2023.04.16.537068

**Authors:** Mélanie Bazin-Gélis, Evangelia Eleftheriou, Coralie Zangarelli, Gaëlle Lelandais, Linda Sperling, Olivier Arnaiz, Mireille Bétermier

**Affiliations:** Université Paris-Saclay, CEA, CNRS, Institute for Integrative Biology of the Cell (I2BC), 91198, Gif-sur-Yvette, France; Present address: Institut Pasteur, Université Paris Cité, Inserm U1223, Innate Immunity Unit, Paris, France

## Abstract

Multinucleate cells are found in many eukaryotes, but how multiple nuclei coordinate their functions is still poorly understood. In the cytoplasm of the ciliate *Paramecium tetraurelia*, two micronuclei (MIC) serving sexual reproduction coexist with a somatic macronucleus (MAC) dedicated to gene expression. During sexual processes, the MAC is progressively destroyed while still ensuring transcription and new MACs develop from copies of the zygotic MIC. Several gene clusters are successively induced and switched off before vegetative growth resumes. Concomitantly, programmed genome rearrangements (PGR) remove transposons and their relics from the new MACs. Development of the new MACs is controlled by the old MAC, since the latter expresses genes involved in PGR, including the *PGM* gene encoding the essential PiggyMac endonuclease that cleaves the ends of eliminated sequences. Using RNA deep sequencing and transcriptome analysis, we show that impairing PGR up-deregulates key PGR genes, together with ∼600 other genes possibly also involved in PGR. Among these genes, 42% are no longer induced when no new MACs are formed, including 180 genes that are co-expressed with *PGM* under all tested conditions. We propose that bi-directional crosstalk between the two coexisting generations of MACs links gene expression to the progression of MAC development.

## INTRODUCTION

Eukaryotic cells generally harbor a single nucleus that serves two major functions: gene expression and transmission of genetic information to post-mitotic daughter cells or post-meiotic gametic cells. However, many examples of multinucleate cells have been documented, such as the muscle fiber of animals, the syncytium of filamentous fungi or the early *Drosophila* embryo (1, 2). In these giant cells, *in situ* hybridization or single-nucleus RNA sequencing have pointed to a heterogeneous transcriptional behavior among individual nuclei or nuclear subtypes occupying different regions of the cytoplasm (3–5). Whether and how the different nuclei of a multinucleate cell coordinate their respective transcriptional activities have been studied in a few cell types (6) but remain overall poorly understood. Ciliates are unicellular eukaryotes, in which germline and somatic functions are partitioned between distinct types of nuclei that coexist in the same cytoplasm (7). Gene transcription takes place in the somatic macronucleus (MAC), a highly polyploid nucleus that is essential for cell survival. The diploid germline micronucleus (MIC), although transcriptionally silent, plays a crucial role in sexual processes: it undergoes meiosis and transmits the germline genome to the zygotic nucleus of the following generation. At each sexual cycle, the old (parental) MAC is destroyed, and in the same cell, two new MACs develop from mitotic copies of the zygotic nucleus. MAC development has been extensively studied in ciliate models (8), including *Paramecium* (9). In *Paramecium tetraurelia*, two sexual processes have been described: conjugation between a pair of reactive partners with complementary mating types, and autogamy, a self-fertilization that takes place in a single cell. At the start of the sexual cycle, MIC meiosis is accompanied by the fragmentation of the old MAC into ∼30 fragments that persist in the cytoplasm and continue to be transcribed throughout new MAC development (10). Gene transcription progressively switches from the old to the new MACs to ensure progeny survival when vegetative growth resumes and old MAC fragments are lost. A large-scale analysis of the *P. tetraurelia* transcriptome during autogamy established that the expression pattern of 17190 genes (out of 41533) follows a developmental program (11), defining six clusters of differentially expressed genes. An early peak of expression takes place during MIC meiosis and old MAC fragmentation. Two successive intermediate and late expression peaks are detected during MAC development, followed by the onset of a late induction cluster. The gene expression program also includes early and late repression clusters.

In *P. tetraurelia*, MAC development involves successive rounds of genome endoduplication to reach the final ploidy (∼1000 to 1600n (12)) of the mature new MAC (10). Most of the Programmed Genome Rearrangements (PGR) start when the new MAC genome has reached a 32n ploidy and extends over several consecutive endoduplication rounds (13). In all, PGR remove at least 30% of germline DNA (reviewed in (9)), reducing the haploid genome complexity from 100 to 150 Mbp in the MIC (14) to 72 to 76 Mbp in the mature MAC (15, 16). Eliminated germline sequences include multiple DNA repeats (transposable elements (TE) and minisatellites) that are removed imprecisely, leading to the formation of variable MAC chromosome ends capped by *de novo* telomere addition and/or to internal deletions (17). In addition, around 45000 Internal Eliminated Sequences (IES), mostly short, non-coding and originating from ancient TEs, are excised from new MAC chromosomes (14, 18) according to a defined temporal program (13). IES ends present a poorly conserved and degenerate consensus sequence, which includes the invariant TA dinucleotide found at IES boundaries with flanking DNA (18, 19). Since 47% of genes are interrupted by at least one IES in the MIC, precise IES excision is necessary to assemble functional genes in the new MAC. Thus, PGR not only shape the structure of MAC chromosomes, but also rearrange the somatic genome to make it suitable for gene expression. Numerous studies have focused on the search for essential actors involved in PGR. A first line of research has unraveled the role of non-coding RNAs (20–23) in guiding the Ezl1 histone-methyl transferase (24), a component of the *Paramecium* Polycomb repressive complex PRC2 (25, 26), to germline-restricted sequences that are eventually eliminated from the developing MAC. Of note, 25% of IESs with stronger nucleotide sequence signals at their ends relative to the general consensus, are excised with high efficiency and precision, independently of non-coding RNAs and heterochromatin marks (13), which suggests that other factors contribute to their recognition. A second line of research has aimed at identifying components of the core IES excision machinery, which are also required for most imprecise DNA elimination (16). A complex of six domesticated PiggyBac transposases, including the catalytically active PiggyMac (Pgm) subunit (27, 28) and five Pgm-like (PgmL) partners (29), has been proposed to form the DNA cleavage complex that introduces DNA double-strand breaks (DSB) at IES ends (30). Once IESs are removed, broken excision sites are repaired through non-homologous end joining (NHEJ): following limited processing of free DNA ends, the NHEJ-specific LigaseIV-Xrcc4 complex stitches together the two IES-flanking sequences (31). In addition to playing its expected role in DSB repair, the specialized Ku70/Ku80c heterodimer appears to be part of the DNA cleavage complex itself: it interacts with Pgm, stabilizes its localization in the developing new MACs and is indispensable for DSB introduction at IES ends (32, 33).

Most of the known proteins that participate in the epigenetic control or catalysis of PGR are encoded by genes from the early or intermediate expression peaks, which are switched off at the end of autogamy in control conditions (11, 34). Previous observations suggested that depleting essential actors of PGR may perturb this gene expression program (32). Indeed, northern blot experiments revealed that *KU80c* and *KU70* mRNAs, which are expressed in the intermediate peak, accumulate continuously until late stages of MAC development in a *PGM* knockdown (KD). Reciprocally, *KU70* and *PGM* mRNAs are up-deregulated in a *KU80c* KD. To generalize these observations, we studied the *P. tetraurelia* transcriptome during autogamy of cells subjected to three independent KDs of individual genes involved in PGR. We report that impairing the normal course of PGR in the new MACs results in up-deregulation of 628 genes, among which we found many known PGR genes, suggesting that the list may contain new important candidate genes for PGR. We also show that subsets of genes from the early and intermediate peaks are no longer expressed when new MACs fail to develop. Based on these findings, we propose that a crosstalk between the two generations of somatic nuclei that coexist in *Paramecium* cells during autogamy establishes a regulatory feedback loop that links the transcription program in the old MAC to the progression of PGR in the new MACs.

## MATERIALS AND METHODS

### *Paramecium* strains and culture conditions

*Paramecium tetraurelia* 51 new (30) or its mutant derivative 51 *nd7-1* (28) were grown at 27°C in a wheat grass infusion medium inoculated with *Klebsiella pneumoniae* (*Kp* medium) and supplemented with 0.8 µg/mL β-sitosterol and 100 µg/mL ampicillin (35).

### Microinjection of the *GFP* reporter transgene

All *GFP* reporter transgenes were constructed from the *PGM-GFP* and *GFP-PGM* constructs published in (28). In plasmid p0349, the *GFP* gene was cloned under the control of *PGM* transcription signals (96 bp upstream of the start codon and 54 bp downstream of the stop codon) (Figure 1A). The wildtype promoter of p0349 was then modified using synthetic DNA fragments (from Integrated DNA Technologies) to give plasmids p0364 and p0418, carrying a mutant or an inverted motif in the *PGM* promoter, respectively. All plasmids were linearized by BsaAI (New England Biolabs) and co-injected into the MAC of vegetative *P. tetraurelia* 51 *nd7-1* with a small amount (1:10 mass ratio) of a linearized *ND7*-complementing plasmid, as described in (28). Transformants were screened for trichocyst discharge (trich+) after 2 to 3 cell divisions. Transformed cells were grown in *Kp* medium for 8-10 vegetative divisions and their genomic DNA was extracted using the NucleoSpin Tissue kit (Macherey-Nagel). Transgene injection levels (copy number per haploid genome, cphg) were determined using real-time qPCR. Oligonucleotide primers used to amplify *PGM* and *GFP*, with *KU80c* as a genomic reference, are listed in Supplementary Table S1. Since there is no endogenous *GFP* gene in *Paramecium*, we used genomic DNA extracted from cells transformed by the *PGM-GFP* fusion as a standard to quantify *GFP* DNA amounts, with a cphg=6 based on Southern blot hybridization (28) and qPCR using *PGM* primers. For all *GFP* transformants, we calculated the apparent cphg from the Cp value provided by real-time qPCR:

**Figure 1.**
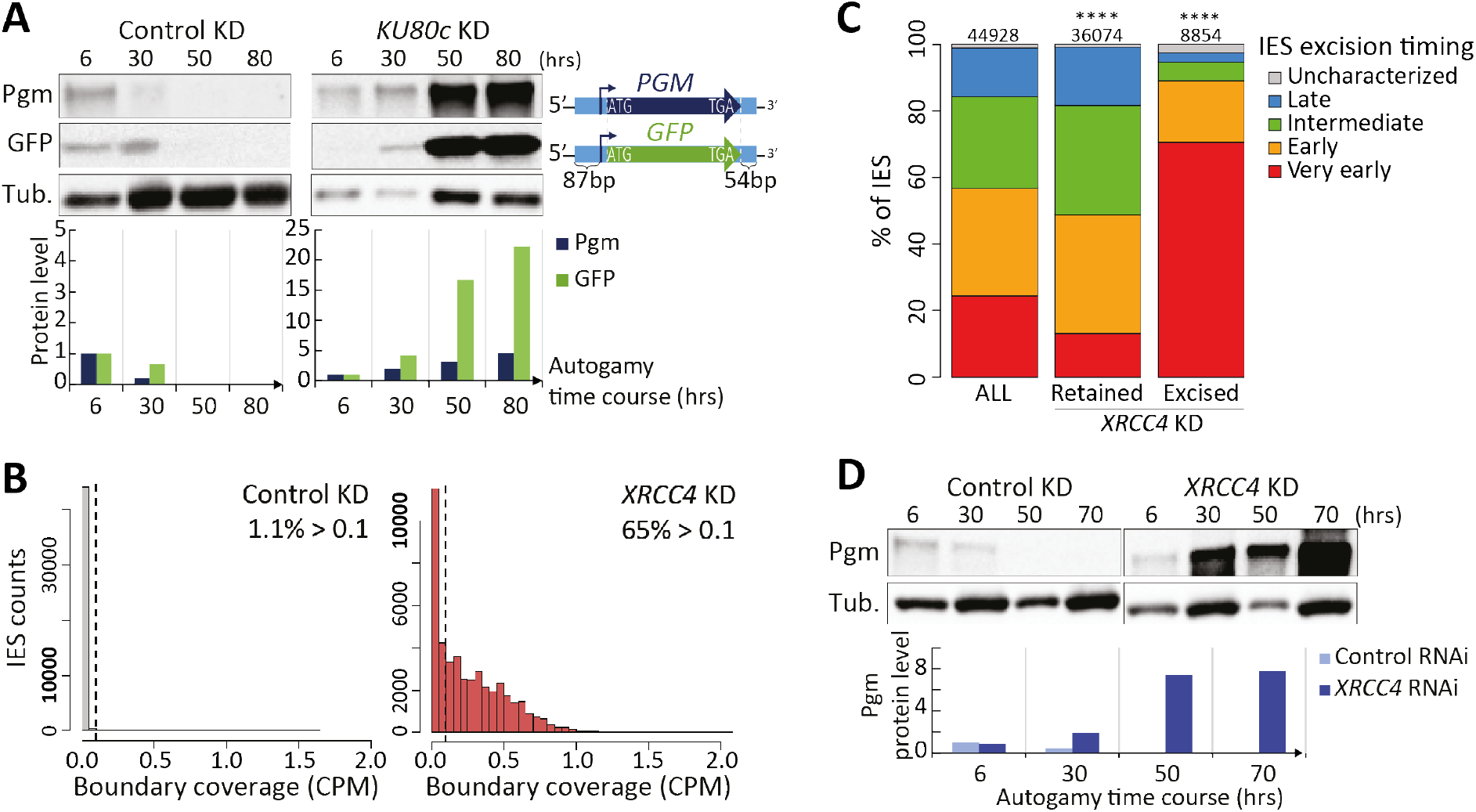
Pgm overexpression in *KU80c* and *XRCC4* KDs. **(A)** Western blot quantification of the levels of endogenous Pgm and transgene-encoded GFP during an autogamy time-course of cells subjected to control or *KU80c* RNAi. The GFP is expressed from a micro-injected transgene (1976 cphg) under the control of *PGM* transcription signals. **(B)** Evidence for IES retention in an *XRCC4* KD. The histograms represent the number of IESs in each 0.05-CPM interval of normalized IES boundary coverage, following control (gray) or *XRCC4* (red) RNAi. The % in each panel indicates the fraction of IESs with a normalized IES boundary coverage >0.1 CPM. In the simulated dataset with full IES retention, 96.2% of IESs had a boundary coverage >0.1 CPM (Supplementary Figure S3). Note that y-axes are not to the same scale in both panels. **(C)** Repartition among the excision timing groups (13) of the IESs with a boundary coverage >0.1 (Retained) or <0.1 CPM (Excised) in *XRCC4* KD compared to all IESs (ALL) (****: p-value <10^-200^, Chi^2^ test). The number of IESs in each category is given above the plots. **(D)** Western blot quantification of endogenous Pgm levels during an autogamy time-course of cells subjected to control or *XRCC4* RNAi. The level of Pgm at T6 in the control RNAi is set to 1 as a reference for all lanes. In panels A and D, time-points are in hours (hrs) following T0. Normalized values (in arbitrary units) are plotted below each western blot.

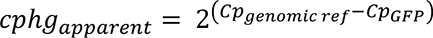

In each experiment a correction factor (f) was calculated from the Cp value obtained with the *GFP* standard:

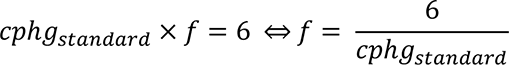

The correction factor f was used to correct the value of the *GFP* injection level (cphg) of transformed cells.

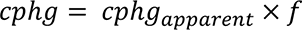

### RNAi-mediated gene knockdown (KD) during autogamy

Double-strand RNA (dsRNA)-induced RNAi was carried out using the feeding procedure (36, 37). Transformed or non-injected *Paramecium* cells were grown in *Kp* medium for 4 vegetative divisions, then transferred to wheat grass infusion medium containing plasmid-free *Escherichia coli* HT115 for 4-5 divisions (38). RNAi was induced as described in (28). Each culture was split and fed for 4-5 divisions with non-induced HT115 containing RNAi plasmids (against target or control genes), then with dsRNA-producing HT115 for 10-15 more divisions before cells starve and start autogamy. Final volumes were 200 mL for medium-scale or 1 L for large-scale experiments. Autogamy was triggered by starvation (39) and the progression of old MAC fragmentation was monitored using DAPI staining of fixed cells (33). In all autogamy time-courses, the T0 time-point is defined as the time when 50% of cells have a fragmented MAC. Three days after autogamy has started, the ability of sexual progeny to develop a functional new MAC was monitored by performing a survival assay as described (28) (Supplementary Figure S1). Control experiments were performed using plasmid p0ND7c, which targets RNAi against the non-essential *ND7* gene (40). The plasmids used for *PGM*, *KU80c*, *XRCC4* and *CtIPa* + *CtIPb (= CtIP)* RNAi were pPBL49HN (27), pL4440-KU80C-2 (32), pXRCC4-R (31), pL4440-PtCtIPa and pL4440-PtCtIPb (41), respectively.

### Cell collection and protein extraction for western blot analysis

50-mL aliquots (∼1 to 2 x 10^5^ cells) were collected from medium-scale cultures at different autogamy time-points. Cells were washed for 20 min in Dryl’s buffer (2 mM sodium citrate, 1 mM NaH_2_PO_4_, 1 mM Na_2_HPO_4_, 1 mM CaCl_2_) before concentration in a final volume of ∼40 µl. 20-µL aliquots were flash-frozen in liquid nitrogen and stored at -80°C. For western blots, 20 µL of frozen cells were lysed by a 3-min incubation at 100°C in 5% boiling SDS with 2x final concentration of Protease Inhibitor Cocktail Set 1 (Merck Chemicals). After centrifugation (5 min 15000g), Laemmli sample buffer (Bio-Rad) with β-mercaptoethanol (354 mM final concentration) was added to the supernatant before a 3-min incubation at 100°C. Electrophoresis was performed in Criterion TGX Stain-free 4-15% precast SDS-Page gels (Bio-Rad) in migration buffer (25 mM Tris-base, 200mM Glycin, 0.1% SDS). Following migration, proteins were electro-transferred to 0.45 μm NC Protran nitrocellulose blotting membranes (Amersham) in transfer buffer (20 mM NaH_2_PO_4_, 20 mM Na_2_HPO_4_, pH 6.7).

Pgm was immuno-detected using α-Pgm 2659-GP guinea pig polyclonal antibodies (1:5000 or 1:1000) (28), GFP using the rabbit polyclonal antibody padg1 (1:2000, ChromoTek), tubulin using the monoclonal α-alpha tubulin TEU435 (1:10000 or 1:5000, kindly provided by A. Aubusson-Fleury and A.M. Tassin) (42). Antibodies were diluted in TBST (50 mM Tris-HCl pH 8, 150 mM NaCl, 0.5% Tween) containing 4% low fat milk powder. Primary antibodies were revealed using species-appropriate HRP-conjugated secondary antibodies (anti-rabbit or anti-mouse IgG (1:5000, Promega), or anti-guinea pig IgG (1:5000, Thermo Scientific)). For each antibody, the signal was acquired using the ChemiDoc Touch Imaging System (Bio-Rad) and quantified with Image Lab software (Bio-Rad) and GFP or Pgm signals were normalized relative to alpha tubulin. Unless otherwise specified, the normalized intensity at T6 was set to 1 as a reference to quantify protein amounts in each time-course.

### RNA sequencing

All RNA preparations used for sequencing were obtained from published autogamy time-course experiments, in which the following genes were knocked down using RNAi: *EZL1* and *ICL7* control (24), *PGM*, *KU80c* & *ND7* control (= ND7_K) (32), *CtIP* (41), *XRCC4* and two independent *ND7* controls (=ND7_X and ND7_L) (31).

RNAseq data from *EZL1* and *ICL7* KDs were previously published (Supplementary Table S2) (25, 43). Poly A RNAs extracted from KU80c, PGM and ND7_K time-courses were sequenced from oriented Illumina TruSeq libraries using a HiSeq sequencer (2 x 100 bp paired end), as described (18). For the XRCC4, CtIP, ND7_X and ND7_L time-courses, RNA sequencing was performed on a NextSeq 500 sequencer (2 x 75 bp paired end), as described (25) (Supplementary Table S2).

### Reference genomes

The sequencing data were mapped on *P. tetraurelia* strain 51 MAC v1 (ptetraurelia_mac_51.fa) and MAC+IES v1 (ptetraurelia_mac_51_with_ies.fa) reference genomes using Hisat2 (v2.1.0, --rna-strandness FR --min-intronlen 20) (11). Gene annotation v2.0 (ptetraurelia_mac_51_annotation_v2.0.gff3) and IES annotation v1 (internal_eliminated_sequence_PGM_ParTIES.pt_51.gff3) were used in this study (11).

### Estimation of IES retention from DNAseq reads in an *XRCC4* KD

For DNA sequencing, developing new MACs were collected at T30 (30 hours after T0) from 500 mL of large-scale cultures of *P. tetraurelia* 51 new grown in *XRCC4* or *ND7* RNAi-inducing media and enriched by centrifugation through a 2.1 M sucrose layer (13, 18). Total DNA from flow cytometry-sorted new MACs (Supplementary Figure S2) was sequenced on a NExtSeq 500 sequencer as described (13) (Supplementary Table S2). To monitor the excision status of each IES, we could not use the standard IES retention score (44) because calculating that score relies on counting *de novo* IES^-^ junctions, which are not formed after DSB introduction in Xrcc4-depleted cells (31). IES retention was therefore evaluated by counting the number of reads mapping on each IES left boundary using ParTIES (MIRET module v1.05 default parameters) (44), normalized by total read coverage per Mbp of the MAC+IES genome (CPM, counts per million) (Supplementary Table S3). As a control, a sequencing dataset with 100% IES retention was simulated using ART_Illumina (v2.5.8 -l 80 -ss HS10 -p -qs 90 -qs2 90 -m 250 -s 50) on the MAC+IES genome, with the same sequencing depth as the XRCC4 KD dataset (Supplementary Figure S3). IES boundary coverage was estimated from the simulated data using the method described above.

### Differential expression (DE) analysis

For each RNAseq dataset, raw fragment counts were determined for each gene using htseq-count (v0.11.2, --stranded=yes --mode=intersection-nonempty) on filtered BAM files (samtools v1.3.1) as described (11). Version 4 of R (45) was used for data analyses and to generate images (ggplot2 v3.3.6; VennDiagram v1.7.3) (46, 47). For the correction of batch effects due to differences in RNA extraction conditions and sequencing protocols (HiSeq or NextSeq), we first validated the ComBat-seq procedure (48) (sva v3.38.0, R package, with no group condition) on technical replicates from the EZL1 and ICL7 time courses that were sequenced using both HiSeq (43) and NextSeq (25) (Supplementary Figure S4, Supplementary Table S4). We then corrected all RNAseq datasets (except for the EZL1 time course) as the first step of our analysis workflow (Supplementary Figure S5), before completing a first round of DESeq2 analysis (49) (DESeq2 v1.30.1, R package). Since biological replicates were available only for control KDs (ND7 and ICL7 time-courses), we used hierarchical clustering (Pearson correlation matrix, stats v4.0.3, R package) and principal component analysis (PCA, FactoMiner v2.4, R package) to group the samples in 4 autogamy stages for each RNAi condition (43) (Part 1 of Supplementary Figure S5). The resulting groups were manually curated based on cytological observations of autogamous cells (Supplementary Figure S6). For the ICL7 time course, the time points that were sequenced both by HiSeq and NextSeq were considered as replicates and collapsed. The samples of all control time-courses grouping in the same autogamy stage were treated together as replicates. For each KD, the time-points that were attributed to the same autogamy stage were considered as replicates in all DE analyses. Following a second DESeq2 round, we validated the final set of 19 replicate groups by hierarchical clustering and PCA analysis (Supplementary Figure S7). The results of DE analysis were extracted using both RNAi conditions and autogamy stages as contrast (Part 2 of Supplementary Figure S5). Genes were considered as deregulated if their fold-change reached at least 2 compared to controls, with an adjusted p-value (padj) below 0.05 (gene lists in Supplementary Tables S4 & S5).

### Identification of a conserved motif

We used STREME 5.4.1 (50) to search for 5 to 20-bp sequence motifs (with a p-value <0.05) that are enriched in the promoters (defined as the 150-bp window upstream of the annotated transcription start site (TSS), see (11)) of the 180 genes that are coregulated with *PGM* (Co-*PGM* genes: intermediate peak genes, up-deregulated in *XRCC4*, *PGM* and *KU80c* KDs and down-regulated in a *CtIP* KD). The search was repeated 5 times against different control sets of the same number of randomly-picked gene promoters. The .meme files corresponding to the first hits of each STREME round were aligned and trimmed to the shortest motif size (18 bp) (trim_motifs, universalmotif v1.12.3, R package). A consensus motif was defined by merging the five motifs (merge_motif, universalmotif v1.12.3, R package). For each round, STREME defined an enrichment factor as the fraction of genes with at least one match to the consensus in their promoter. The significance threshold for further screening was based on the mean of the 5 enrichment factors.

### Analysis of the motif in gene promoters

To identify all the genes that carry the motif in their promoter, we used the FIMO 5.4.1 software (51) with default settings. We first extracted the data obtained for the set of 180 Co-*PGM* genes and defined a cut-off p-value that matches the significance threshold calculated from STREME enrichment factors (defined above). We then applied the resulting cut-off to the full list of motifs identified by FIMO. To compare the position of the motifs in their respective promoter, we considered the position of the first 5’ nucleotide of the motif relative to the TSS and counted the number of motifs found in successive 5-bp windows within gene promoters.

## RESULTS

### Inhibition of Programmed Genome Rearrangements (PGR) triggers overexpression of the Pgm endonuclease

Following up on previous experiments (32), we confirmed on western blots that the observed up-deregulation of *PGM* mRNA in a *KU80c* KD results in accumulation of Pgm at late developmental stages (Figure 1A, Supplementary Figure S1A for survival test). To investigate whether the deregulation takes place at the transcriptional level, a *GFP* reporter transgene expressed under the control of *PGM* transcription start and termination signals was micro-injected into the MAC of vegetative *P. tetraurelia* cells. GFP levels were monitored on western blots after the onset of autogamy (Figure 1A). We observed that accumulation of GFP in a *KU80c* KD parallels that of endogenous Pgm, indicating that gene deregulation mostly takes place in the old MAC. We cannot, however, ascertain whether the accumulation of GFP results from transcriptional or post-transcriptional deregulation, since the mRNA produced from the transgene carries both the 5’ and 3’ UTRs of *PGM*.

Pgm and Ku70/Ku80c are essential for DNA cleavage upon initiation of IES excision (27, 32). As a consequence, RNAi-mediated silencing of either *PGM* or *KU80c* causes the retention of all IESs (18, 33). To examine whether interfering with DSB repair after IES end cleavage, a step that is necessary for genome assembly in the new MACs, would also up-deregulate *PGM*, we knocked down *XRCC4*. Previous molecular analyses performed on a handful of IESs established that DSBs introduced at IES ends in an *XRCC4* KD are not repaired, resulting in the persistence of broken ends and a failure to assemble IES-junctions (31). At the genome-wide level, high-throughput DNA sequencing from purified new MACs revealed that 65% of IESs were retained at T30 in an *XRCC4* KD, in contrast to a control KD (Figure 1B). Of note, this set of retained IESs is depleted in the earliest excised IESs during autogamy (very early IESs) (13) (Figure 1C). Reciprocally, very early excised IESs are overrepresented among the 8854 IESs exhibiting little or no retention. This shows that inhibiting DSB repair after the first IESs are cut out of chromosomes impinges on the normal progression of PGR and results in late IES retention. As shown on western blots, we observed a strong increase in Pgm levels in an *XRCC4* KD relative to control conditions (Figure 1D). We conclude that knocking down *XRCC4* also deregulates *PGM* expression during MAC development, even if IES excision is not affected through the same mechanism as in a *KU80c* KD.

### Genome-wide identification of up-deregulated autogamy genes in *PGM*, *KU80c* and *XRCC4* KDs

*PGM* belongs to a cluster of 2037 differentially expressed genes that are transiently induced during autogamy, clustering in the intermediate expression peak, when PGR take place in the developing MACs (11). To gain broader insight into how many genes are up-deregulated in the three KDs that we used in the present study to alter the normal course of PGR, we collected RNA sequencing data from independent autogamy time-course experiments performed on cells subjected to *KU80c*, *PGM* (32) or *XRCC4* KDs (31) and compared them with four independent control time-courses. We applied principal component analysis and hierarchical clustering to the normalized datasets to group the samples in four developmental stages (VEGETATIVE, EARLY, INTERMEDIATE and LATE) (Supplementary Figure S7). These stages coincide with the time-windows when genes from the early, intermediate and late expression peaks reach their maximal expression level (11). To define the sets of genes that are deregulated when PGR are impaired, we focused on the LATE stage, which yielded the largest number of deregulated genes in each KD relative to the controls (Supplementary Table S5). We found that many genes are significantly up-deregulated at the LATE stage in a *PGM*, *KU80c* or *XRCC4* KD (Figure 2A). Of note, fewer up-deregulated genes were identified in Ku80c-than Pgm-depleted cells, whereas *PGM* and *KU80c* KDs cause similar retention of all IESs (18, 33). This is likely due to faster progression of autogamy in the *PGM* RNAi experiment (Supplementary Figure S6) leading to a more contrasted difference in transcript levels relative to controls at the LATE stage. We defined a set of 628 genes, referred to as UP PKX genes, that are up-deregulated in all three KDs (Figure 2A). Almost half of the 628 UP PKX genes belong to the early (15%) or intermediate (37%) expression peaks (11) (Figure 2B), an enrichment that can also be detected in each individual KD (Supplementary Figure S8A-D). We also noted a bias in the germline structure of the LATE up-deregulated genes, 86% being devoid of any IES compared with 53% for all genes (Figure 2C) and 55% on average for random sets of 628 genes with similar proportions of each expression cluster (Supplementary Figure S8E-H). The 235 LATE up-deregulated genes from the intermediate peak, which represent only ∼12% of the expression cluster, include almost all of the genes encoding known components of the DNA cleavage complex (Supplementary Table S6): *PGML1, PGML2, PGML3a, PGML4a&b, PGML5a&b* (29) and *KU70a* (32). Only *PGM* (27) and *KU80c* (32) are missing since, even if up-deregulated in the other two KDs, they are not overexpressed in their own respective KD. Our results suggest that mRNAs from genes involved in PGR accumulate (or persist) at LATE stages in *PGM*, *KU80c* or *XRCC4* KDs as a response to defective IES excision in the new MAC.

**Figure 2.**
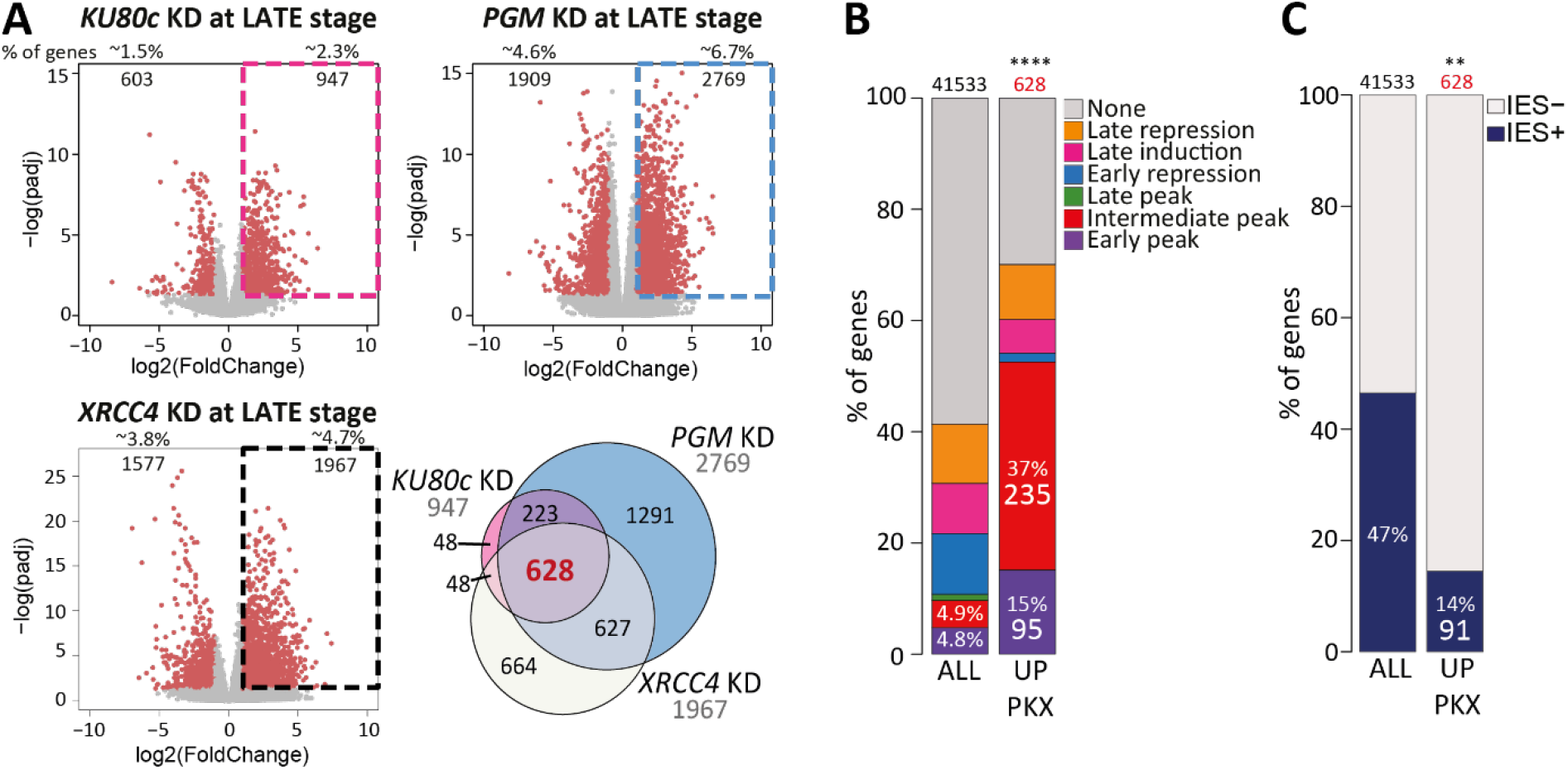
Up-deregulated genes in *PGM*, *KU80c* and *XRCC4* KDs at the LATE autogamy stage. **(A)** Volcano plots showing differentially expressed genes at the LATE stage in *PGM* (top left), *KU80c* (top right) or *XRCC4* (bottom left) KDs relative to the controls. The most deregulated genes are indicated (in red), with their number and percentage relative to all 41533 genes. Up-deregulated genes are boxed on each plot (padj: DESeq2 adjusted p-value). The Venn diagram shows the intersection of the sets of genes that are up-deregulated at LATE stages in all three KDs. **(B)** Proportions of gene from each previously identified expression cluster (11) among the different sets of deregulated genes. ALL: all annotated genes; UP PKX: up-deregulated genes at LATE stage in *PGM*, *KU80c* and *XRCC4* KDs (****: p-value <10^-200^, Chi^2^test). **(C)** Percentage of genes without IES (IES-) or carrying at least one IES (IES+) among the two categories described in B (**: p-value <10^-20^, Chi^2^test).

### The developing new MACs control the transcriptional program throughout MAC development

The shut-off of 330 genes from the early and intermediate expression peaks at the LATE stage of autogamy (95 and 235 genes respectively) seems to critically depend upon the correct completion of PGR in the new developing MACs (Figure 2B), a similar behavior to that of endogenous *PGM* and its reporter *GFP* transgene (Figure 1A and B). This suggests that gene transcription in the old MAC, which drives the production of PGR factors imported in the new MACs, is turned off at the LATE autogamy stage by some signal(s) coming from the new developing MACs, in a regulatory feedback loop.

To investigate whether the new MACs also control gene transcription at earlier stages of autogamy, we examined the production of Pgm in cells depleted in CtIP, an essential protein involved in MIC meiosis (41). In a *CtIP* KD, the formation of the zygotic nucleus is inhibited and no new MAC is formed (Figure 3A, Supplementary Figure S6). In contrast to control conditions, we detected no expression of endogenous Pgm in a *CtIP* KD, whichever autogamy stage was examined (Figure 3B). Following micro-injection of the reporter *GFP* transgene into the old MAC, we observed only a background GFP signal in a *CtIP* KD (Figure 3C and Supplementary Figure S9). Our data show that *PGM* expression from the old MAC is abolished when no new MAC develops after the onset of autogamy. We then extended our study to the whole transcriptome of *CtIP*-silenced cells (Supplementary Table S2) and performing DE analyses. To identify genes behaving as *PGM*, we searched for genes with a reduced expression level in a *CtIP* KD relative to controls, at the EARLY and INTERMEDIATE autogamy stages (Figure 3D). We found 2365 significantly down-deregulated genes at either of these stages, with an enrichment in genes from the early or intermediate peaks and the late induction cluster (Figure 3E and Supplementary Figure S8I) but no significant bias in IES content (Figure 3F and Supplementary Figure S8J). Of note, 263 down-deregulated genes in a *CtIP* KD at EARLY or INTERMEDIATE stages are also up-deregulated in *PGM*, *KU80c* and *XRCC4* KDs at LATE stages (Figure 3D). Among these, 180 belong to the intermediate peak and thus follow the same expression pattern as *PGM* under all the conditions that we tested in this study. Our data suggest that these 180 genes, hereafter designated as Co-*PGM* genes, are co-regulated with *PGM*. Their expression is turned on in the old MAC at the EARLY/INTERMEDIATE stages and remains on until PGR are completed. We propose that regulatory signals originating from the developing new MACs contribute to controlling this expression pattern.

**Figure 3.**
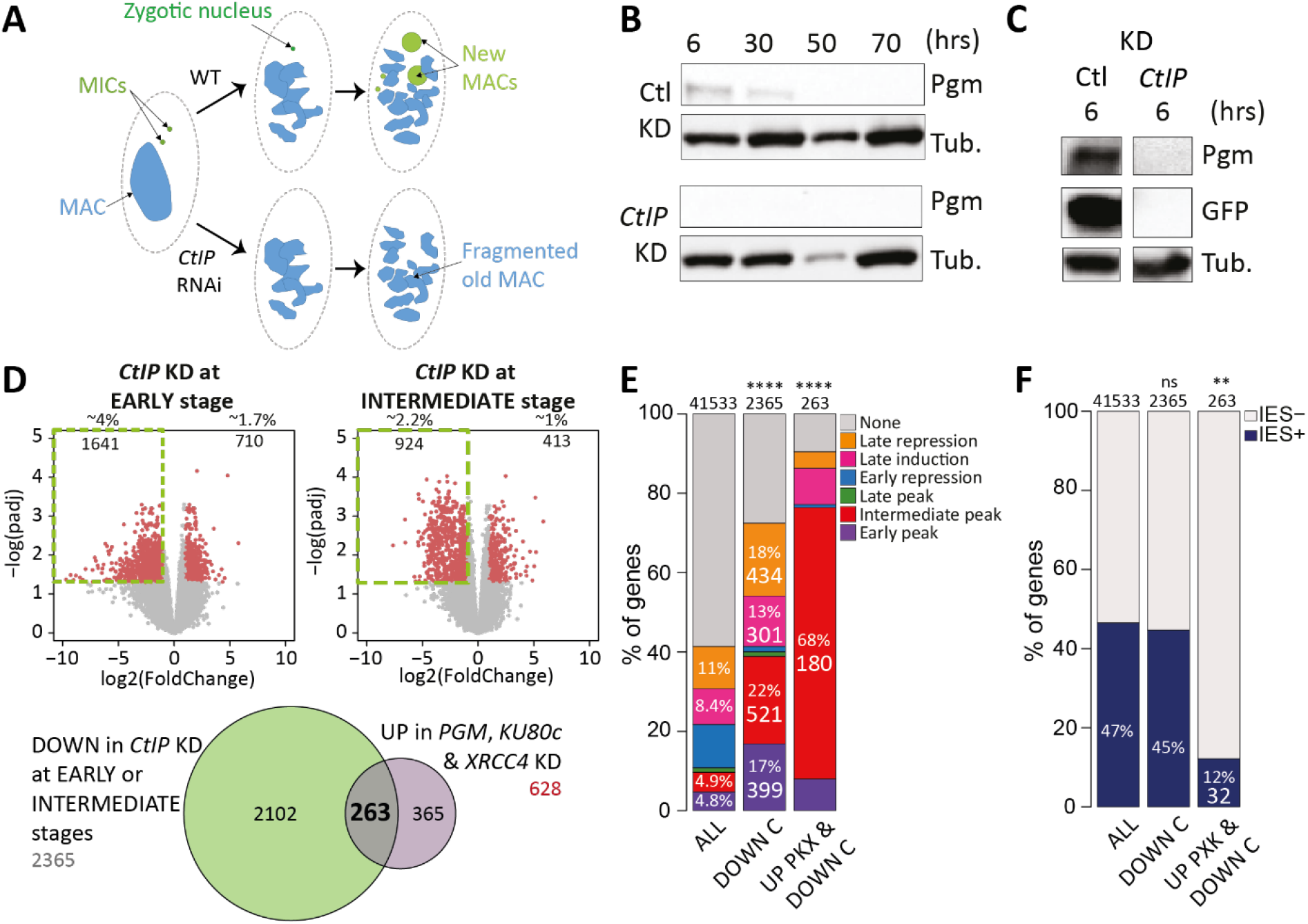
Down-deregulated genes in a *CtIP* KD at the EARLY or INTERMEDIATE stages. **(A)** *CtIP* RNAi inhibits the formation of the zygotic nucleus and the development of new MACs, while old MAC fragmentation still takes place (41). (B) Western blots showing Pgm levels during autogamy time-courses in control (Ctl) or *CtIP* KDs. Time-points are in hours (hrs) following T0. The α-tubulin (Tub.) was used as a loading control **(C)** Comparison of the endogenous Pgm and GFP signals on western blots at T6 for cells harboring the *GFP* reporter transgene (416 cphg), following control or *CtIP* RNAi. The whole membrane is displayed in Supplementary Figure S9A. **(D)** Volcano plots showing differentially expressed genes at the EARLY (left) or INTERMEDIATE (right) stages in a *CtIP* KD relative to the controls. Significantly deregulated genes are indicated (in red), with their number and percentage relative to all 41533 genes. Down-deregulated genes are boxed. The Venn diagram shows the intersection of the sets of genes that are down-deregulated in a *CtIP* KD at EARLY or INTERMEDIATE stages, and up-deregulated at LATE stages in all three conditions displayed in Figure 2. **(E)** Proportions of genes from each previously identified expression cluster (11) among the different sets of deregulated genes. ALL: all annotated genes; DOWN C: down-deregulated genes at EARLY or INTERMEDIATE stages in *CtIP* KD; UP PKX & DOWN C: intersection of the previously described DOWN C genes and the up-deregulated genes at LATE stage in *PGM*, *KU80c* and *XRCC4* KDs (****: p-value <10^-200^, Chi^2^test). **(F)** Percentage of genes without IES (IES-) or carrying at least one IES (IES+) among the three categories described in E (**: p-value <10^-20^; ns: p-value >0.05 Chi² test).

### A conserved regulatory motif is enriched in the promoter of Co-*PGM* genes

To get further insight into the mechanisms that may underlie the transcriptional control of Co-*PGM* genes, we used STREME to search for conserved nucleotide sequence motifs in their promoters. We found a significantly enriched 18-bp non-palindromic consensus motif (5’-AAAATCMTTdWAATAWTT-3’, where M: A or C; d: A, G or T; W: A or T) (Figure 4A), present in either orientation in 98 out of the 180 promoters of Co-*PGM* genes (54%; Figure 4B). At the whole-genome level, a total of 3704 genes carried one (94% genes) or more (6% genes) copies of the motif in their promoter (Supplementary Table S7). Apart from the 98 genes of the Co-*PGM* subset, the motif was also found in the promoter of 3606 other genes (Figure 4B), including 261 genes from the intermediate expression peak (InterP*) that were not deregulated at least in one of the four KDs that we tested in this study (Figure 4C). No particular orientation bias was noted for the motif (Figure 4D). However, the position of the motif upstream of the TSS was markedly different for the 98 Co-*PGM* genes, with a significantly higher density in the -60/-50-bp region (Figure 4E).

**Figure 4.**
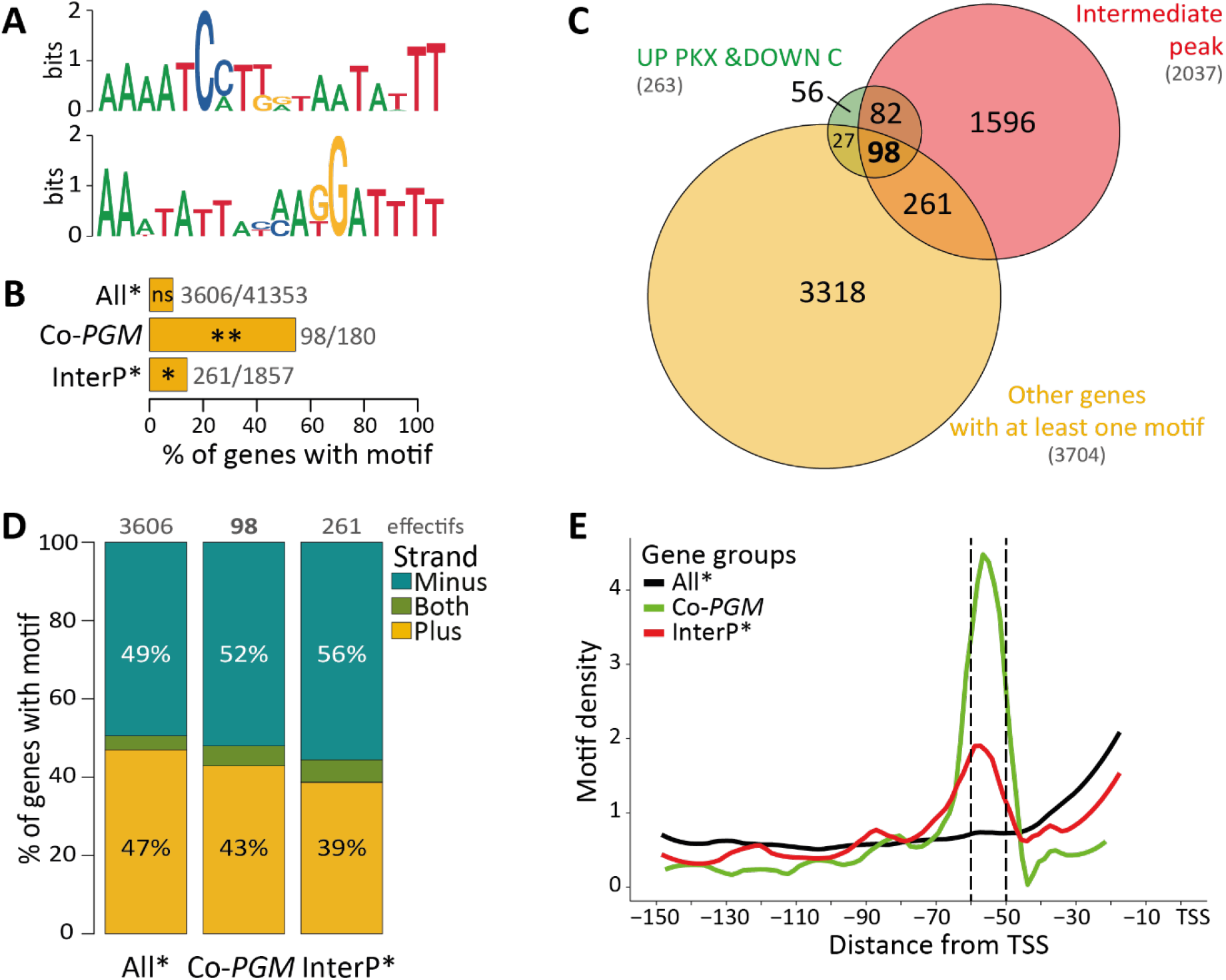
Identification and characterization of an enriched motif in the promoter of Co-*PGM* genes. **(A)** Logo of the 18-bp non-palindromic merged motif (top) and its reverse complement (bottom) identified by STREME in the 150bp upstream of gene TSSs (Transcription Start Sites). **(B)** The enrichment percentage of genes with at least one motif in their promoter is shown relative to all 41533 genes (All*: all genes except Co-*PGM* genes, InterP*: all intermediate peak genes excluding Co-*PGM* genes). **: p-value <10^-20^; *: p-value <10^-10^; ns: p-value >0.05 (Chi^2^ test). **(C)** Venn diagram showing the intersection of the set of genes that are up-deregulated genes in *PGM*, *KU80c* and *XRCC4* KD at LATE stage and down de-regulated in *CtIP* KD at EARLY or INTERMEDIATE stage (green), with the intermediate peak expression cluster (red) and all genes carrying at least one motif in their promoter (yellow). **(D)** Orientation of the motif in gene promoters for the three gene categories described in B. (Minus: all motifs are on the minus strand; Plus: all motifs are on the plus strand; Both: the promoter carries at least two motifs in different orientations). **(E)** Motif density as a function of the distance of the 5’ nucleotide from the TSS, for the three gene categories described in B.

To address the biological significance of the conserved motif, we focused on the *PGM* gene itself. Indeed, a copy of the motif is present 55 bp upstream of the *PGM* TSS in *P. tetraurelia* and is conserved at the same position in other *Paramecium* species (Figure 5A; Supplementary Figure S10). We mutagenized the motif upstream of our *GFP* reporter transgene and, following micro-injection into the vegetative MAC, we followed on western blots the production of GFP during autogamy of transformed cells harboring either the wild-type or the mutant construct (Figure 5B, Supplementary Figure S11). No GFP production was detected at any stage from the transgene carrying the mutant motif, in the control or the *KU80c* KD, suggesting that the motif is essential for gene transcription. We then tested the effect of changing the orientation of the motif by replacing it with its reverse complement upstream of the *GFP* reporter gene (Figure 5A). In the control, the pattern of GFP production was unchanged during autogamy, with a maximum at T7 and a decrease at late stages, even though protein levels were overall lower than with the wild-type construct (Figure 5C, left panels). In the *KU80c* KD, GFP expression from the promoter carrying the inverted motif was up-deregulated at late autogamy stages, similar to what we observed for the wild-type promoter (Figure 5C, right panel). We conclude from these experiments that the conserved motif is required for *PGM* gene expression independently of its orientation.

**Figure 5.**
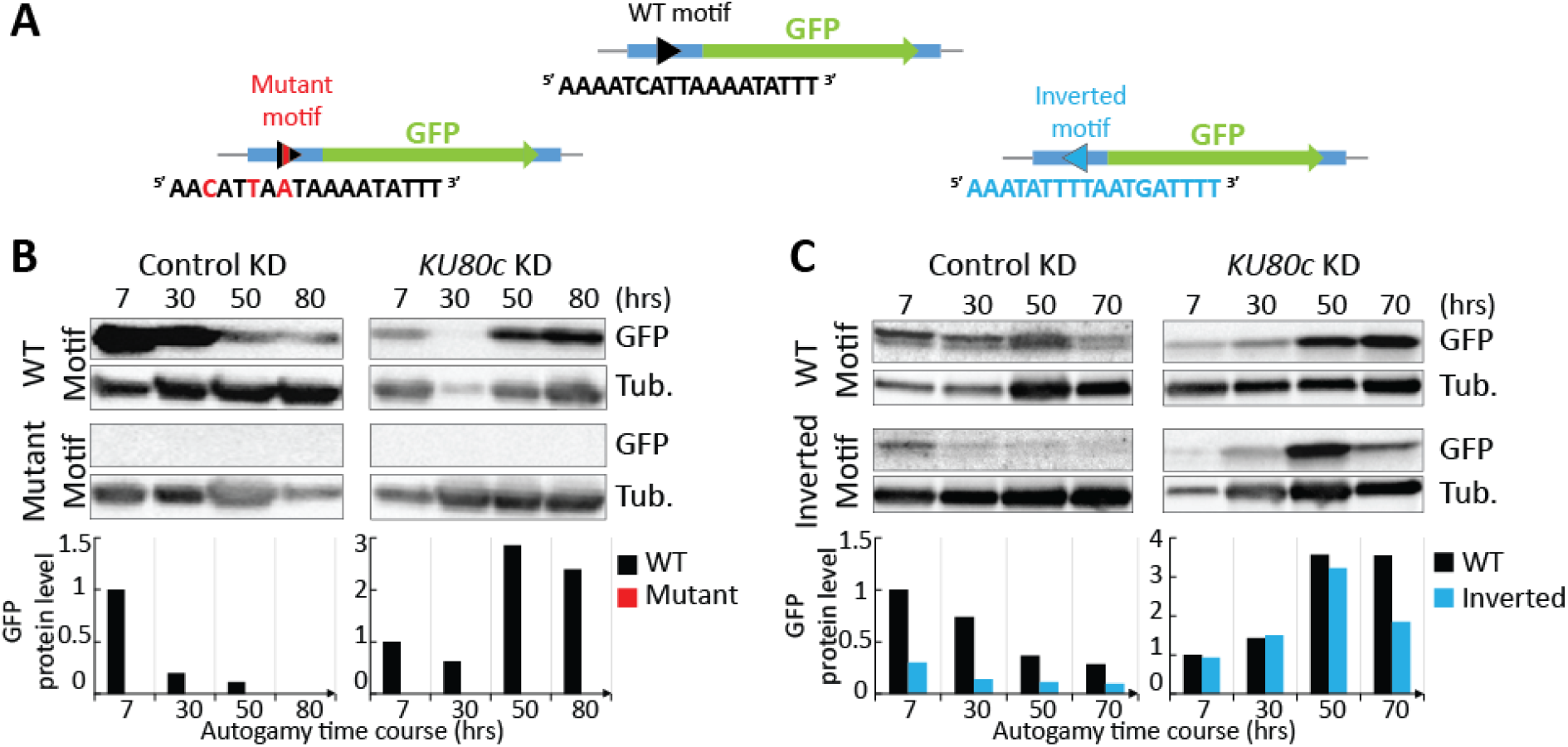
Functional analysis of the motif using the *GFP* reporter transgene. **(A)** Schematic representation of the *GFP* reporter transgenes (see Figure 1) carrying either the wild-type (WT), mutant or inverted motif in their promoter (see sequences below each diagram). In B and C, time-points are in hours (hrs) following T0. Normalized values relative to the α-tubulin (Tub.) signal are plotted below each western blot. In each panel, the level of protein produced at T7 from the WT transgene is arbitrarily set to 1 and used as a reference. **(B)** Western blot quantification of the levels of endogenous Pgm and the GFP expressed from the reporter constructs (WT: 695 cphg, Mutant: 2524 cphg), during an autogamy time-course of cells subjected to control or *KU80c* RNAi. **(C)** Western blot quantification of the levels of endogenous Pgm and the GFP expressed from the reporter constructs (WT: 204 cphg, Inverted: 188 cphg), during an autogamy time-course of cells subjected to control or *KU80c* RNAi.

## DISCUSSION

*Paramecium* constitutes an intriguing example of a multinucleate cell, in which structurally distinct nuclei fulfill different functions. In this study, we have focused on a particular stage of the sexual cycle, during which two consecutive generations of somatic nuclei coexist in the same cell. At this stage, the old (parental) MAC ensures most gene expression, while the new (zygotic) MACs differentiate during a process that involves massive PGR in their genome, before taking over transcription in the progeny. Sequential waves of gene induction and repression were described during autogamy (11). However, what controls the progression of the transcription program during the sexual cycle is still poorly understood. We show here that abolishing the formation of new MACs (*CtIP* KD) or perturbing the normal course of PGR in the new MACs (*PGM*, *KU80c* or *XRCC4* KDs) induce significant variations of the *P. tetraurelia* transcriptome during autogamy. Our results provide evidence that the new MACs control, at two different levels, the progression of the developmental transcription program in the old MAC.

### Developing new MACs control gene expression during autogamy

The first level of *trans*-nuclear control takes place at the EARLY and INTERMEDIATE stages of autogamy. Several published functional complementation experiments using microinjected transgenes confirmed that, consistent with a former study performed during conjugation (10), the old MAC contributes most of the mRNAs of known PGR genes during autogamy, which is sufficient to ensure that functional new MACs develop in the offspring (25, 26, 28, 33, 43, 52). In a *CtIP* KD, the failure to form a zygotic nucleus completely abolishes the development of new MACs without impinging on the fragmentation of the old MAC (41), from which most mRNAs are produced during autogamy. Our transcriptome analysis indicates that the absence of new developing MACs reduces the steady-state mRNA levels of 2365 genes at the EARLY and INTERMEDIATE stages (Figure 3), half of which are specifically induced during autogamy under control conditions (11). These include a fraction of genes from the early and intermediate peaks (20% and 26%, respectively) and the late induction cluster (8%).

The second level of control exerted by the new MACs depends on the outcome of PGR. Our work extends previous observations that impeding PGR in the new MACs triggers gene up-deregulation at LATE autogamy stages (32). Using three independent KDs interfering with PGR at different steps, we report that 628 so-called UP PKX genes are up-deregulated at the LATE autogamy stage in *PGM, KU80c* and *XRCC4* KDs, instead of being switched off as in the control (Supplementary Table S6). We did not include EZL1 RNA-seq data in our DE analysis because, as part of the PRC2 complex, Ezl1 is expected to repress gene expression through heterochromatin formation, as established in other species (53, 54). We nevertheless compared the set of UP PKX genes with the list of LATE up-deregulated genes in an *EZL1* KD (Supplementary Table S4) and found a large overlap between the two sets of genes, with UP PKX genes representing ∼20% of up-deregulated genes in an *EZL1* KD (Supplementary Figure S12). Thus, two effects of *EZL1* KD on gene expression may be distinguished in our analyses: LATE overexpression of the set of UP PKX genes seems to be a consequence of PGR inhibition (24, 43), whereas up-deregulation of other genes is probably due to loss of Ezl1-mediated chromatin modification, independently of PGR.

Under control conditions, 15% of UP PKX genes are expressed in the early peak and 37% in the intermediate peak, which corresponds to the expression pattern of most known PGR genes (11). In addition to *PGM* and *KU80c*, we found that 8 UP PKX genes encode essential components of the DNA cleavage complex (*PGML1, PGML2, PGML3a, PGML4a&b, PGML5a&b* and *KU70a*) (29, 32). Ten more UP PKX genes are involved in RNA-mediated targeting of eliminated DNA (*NOWA1* & *NOWA2* (55), *TFIIS4* (23), *PDSG2* (56)), the control of accessibility of DNA sequences to the cleavage complex through chromatin remodeling (*SPT16.1* (52), *ISWI1* (57)) or take part in PGR through still unclear mechanisms (*DIE5a* (58), *mtGa*&*b* (59), *SPO11* (60)) (Supplementary Figure S12). Based on these observations, we propose that the set of 628 UP PKX genes may include new candidates, whose function in PGR should be explored further.

Since a *GFP* reporter transgene expressed from the old MAC under the control of *PGM* transcription signals shows a similar expression pattern to endogenous *PGM*, we conclude that LATE up-deregulation applies to old MAC transcripts. However, the observation that 86% of UP PKX genes are IES-free (Figure 2C), while 74% of down-deregulated genes carry at least one IES (Supplementary Figure S8E-H), suggests that deregulation of mRNA production in *PGM*, *KU80c* or *XRCC4* KDs also takes place in the new MACs at the LATE stage. The contribution of new MACs to LATE transcription is supported by the detection of increasing amounts of IES+ transcripts at this stage in the three KDs (Supplementary Figure S13). However, the underrepresentation of IES-carrying genes among UP PKX genes may be explained by the selective degradation of IES+ transcripts through Nonsense-Mediated Decay (NMD), because IES retention frequently introduces premature termination codons in mRNAs (18). Consistent with a previous report (61), we indeed found that *PTIWI10*, a gene involved in non-coding RNA-mediated control of IES excision, is down-deregulated at LATE stages in *PGM, KU80c* and *XRCC4* KDs (Supplementary Table S6). *PTIWI10* is interrupted by an NMD-visible IES in the non-rearranged genome and was shown to be specifically expressed from the new MACs (61). In *PGM* and *KU80c* KDs, retention of the IES may trigger NMD-mediated degradation of *PTIWI10* mRNA. This IES, which belongs to the very early excised class, is not retained in an *XRCC4* KD: we propose that its ends are cleaved by the DNA cleavage complex but not repaired, leading to the persistence of an intragenic DSB that abolishes the production of full-length *PTIWI10* mRNA. Taken together, our observations suggest that LATE up-deregulation also takes place in the new MACs but, as a consequence of NMD or unrepaired chromosomal DSBs at excision sites, IES-containing genes expressed from the new MACs may have been overlooked in our search for UP PKX genes.

### Bi-directional crosstalk between two generations of somatic nuclei in a single cell

Several lines of evidence have established that the old MAC controls new MAC development through several pathways. First, maternal non-coding transcripts produced by the old MAC serve as pairing templates for the selection of MIC-restricted small non-coding RNAs that are transferred to the new MACs (22), in which they drive the recognition of eliminated sequences through Ezl1-mediated formation of heterochromatin (25, 26). Second, as discussed above, coding transcription in the old MAC ensures the synthesis of key proteins that carry out PGR in the new MACs.

Here, we provide evidence that, reciprocally, the new developing MACs control the onset and progression of the gene expression program in the old MAC throughout the successive stages of autogamy (Figure 6). The mechanistic details of this *trans*-nuclear control are still unclear and may involve different pathways. We have nevertheless identified a group of 180 genes that are co-regulated with *PGM* under all tested conditions (Co-*PGM* genes, Figure 3E): these genes are specifically induced in the intermediate expression peak, their induction depends on the presence of new developing MACs in the cell and their mRNA levels drop down after the successful completion of PGR. Future studies will unravel the mechanism(s) that underlie the control of gene expression by the new MACs. One attractive hypothesis postulates that the production of activating factors is triggered by the developing new MACs to turn on gene transcription in the old MAC at EARLY and INTERMEDIATE stages. Our finding that 98 out of 180 Co-*PGM* genes carry a conserved motif in their promoter, like *PGM* itself, suggests that genes from this subset are activated through the sequence-specific binding of a transcription factor to the motif, while the expression of Co-*PGM* genes lacking the motif is likely governed by a different regulatory pathway. The observed LATE up-deregulation of Co-*PGM* genes when PGR do not proceed normally may indicate that the activator(s) is expressed from a non-rearranged locus in the new MACs and/or that a repressor produced after PGR are completed is necessary to switch off gene expression.

**Figure 6.**
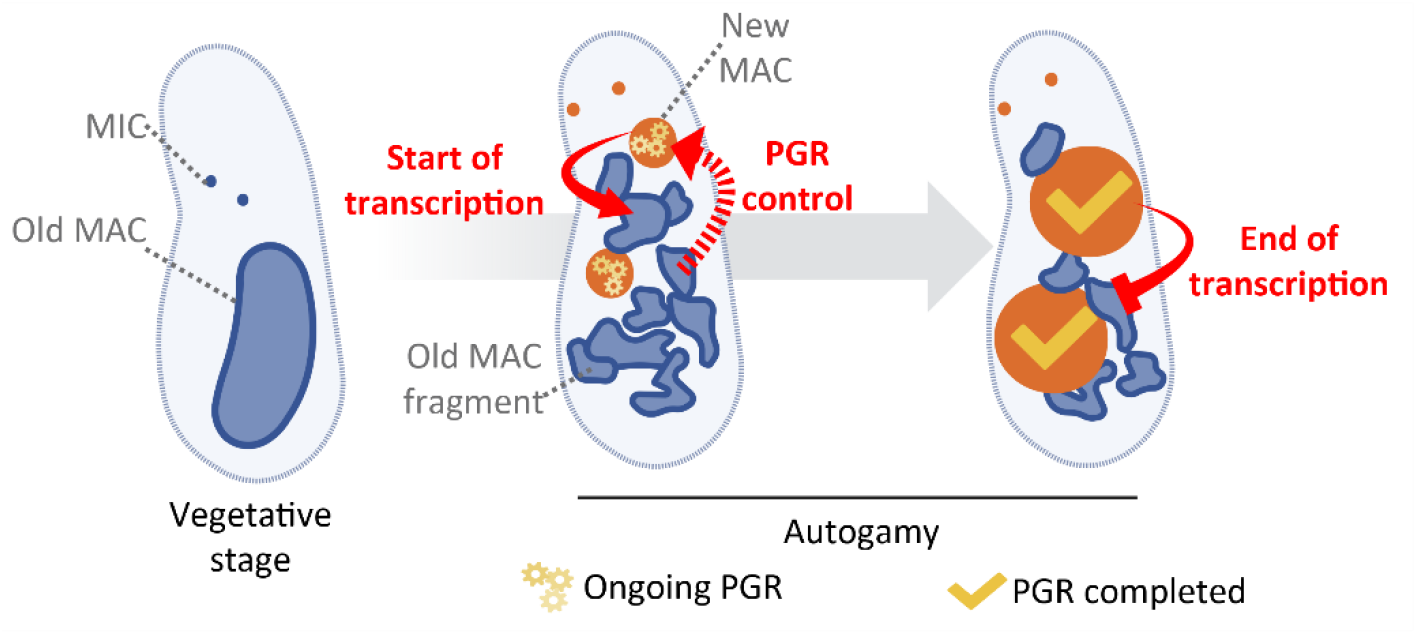
Bi-directional crosstalk between the old and new somatic MACs in *Paramecium*. Upon starvation, vegetative cells (left) become committed to starting their sexual cycle. During autogamy, the MICs undergo meiosis and eventually give rise to the new MICs and MACs (orange), while the parental MAC (blue) is fragmented into ∼30 fragments. Most mRNAs are produced by the old MAC fragments, while programmed genome rearrangements (PGR) take place in the new MACs. We propose that, at EARLY and INTERMEDIATE stages (middle), the new MACs transmit an inducing signal to old MAC fragments (full arrow) to start the transcription of development-specific genes. This drives the production of proteins that carry out PGR in the new MACs. The old MAC also produces constitutive maternal ncRNAs, which control the determination of sequences that will be eliminated from the new MAC (22). The dotted arrow represents the contribution of the old MAC to the control of PGR in the new developing MACs. In turn, we propose that, at the LATE stage (right), a signal from the new MACs arrests the expression of a subset of genes when PGR are completed. (Created with BioRender.com).

The existence of bi-directional cross-communication between the two successive generations of somatic nuclei throughout the *Paramecium* sexual cycle is reminiscent of the mother-embryo crosstalk that has been reported during mammalian development (see (62) for a review). In both systems, the parental and zygotic somatic lines appear to mutually control each other through the production of coding and non-coding RNAs, proteins and other regulatory factors. A notable difference is that inter-generational exchanges in *Paramecium* are intra-cellular: how a continuous dialogue is established at the molecular level between the old and the new developing MACs will require further investigation. We propose that inter-nuclear crosstalk allows gene transcription in the old MAC and genome rearrangements in the new MACs to proceed in a concerted manner to give rise to viable sexual progeny.

## DATA AVAILABILITY

Genes used in this study are accessible in ParameciumDB (https://paramecium.i2bc.paris-saclay.fr/)) as follows: *CtIPa* (PTET.51.1.G0650078), *CtIPb* (PTET.51.1.G0980137), *EZL1* (PTET.51.1.G1740049), *ICL7a* (PTET.51.1.G0700039), *KU80c* (PTET.51.1.G1140146), *ND7* (PTET.51.1.G0050374), *PGM* (PTET.51.1.G0490162) and *XRCC4* (PTET.51.1.G1110086) (Arnaiz et al. 2020).

The sequencing data generated for this study have been submitted to the ENA database (https://www.ebi.ac.uk/ena/browser/home) under accession number PRJEB60900. Plasmid maps, raw images, statistical data, scripts (Supplemental Codes) and numerical data used to produce all diagrams have been deposited at Zenodo (https://doi.org/10.5281/zenodo.7822015).

## ACKNOWLEDGEMENTS

We would like to thank Sandra Duharcourt and Eric Meyer for fruitful discussions, sharing data ahead of publication and providing helpful advice and support as members of MBG’s thesis advisory committee. We are grateful to Anne Aubusson-Fleury and Anne-Marie Tassin for the generous gift of TEU435 antibodies. We thank Joël Acker, Julien Bischerour and Valerio Vitali for critical reading of the manuscript, all members of the Bétermier lab for stimulating discussions, Pascaline Tirand and Cindy Mathon for technical assistance and Tim Kamara and Camille Poitrenaud for help in plasmid construction during their short-term internships.

