## Supplementary Information for "Inter-generational nuclear crosstalk links the control of gene expression to programmed genome rearrangements during the *Paramecium* sexual cycle"

**Supplementary Table S1. Oligonucleotides used as qPCR primers for cphg calculation**

| Amplified regions | ID | Sequences |
|---|---|---|
| *PGM* | OMB433 | GCAGTGGGAACTATTAGACATAACAGAG |
| | OMB434 | TGGAAATACATGAGCATAAGTTCATTTG |
| *KU80c* | OMB1176 | AGCTCCTGATGGCAAGACAT |
| | OMB1177 | AGTCTGTCCACAACCAGCTG |
| *GFP* | OMB1255 | GCCAACACTTGTCACTACTTTAA |
| | OMB1256 | ACTTCAGCTCTTGTCTTGTAGT |

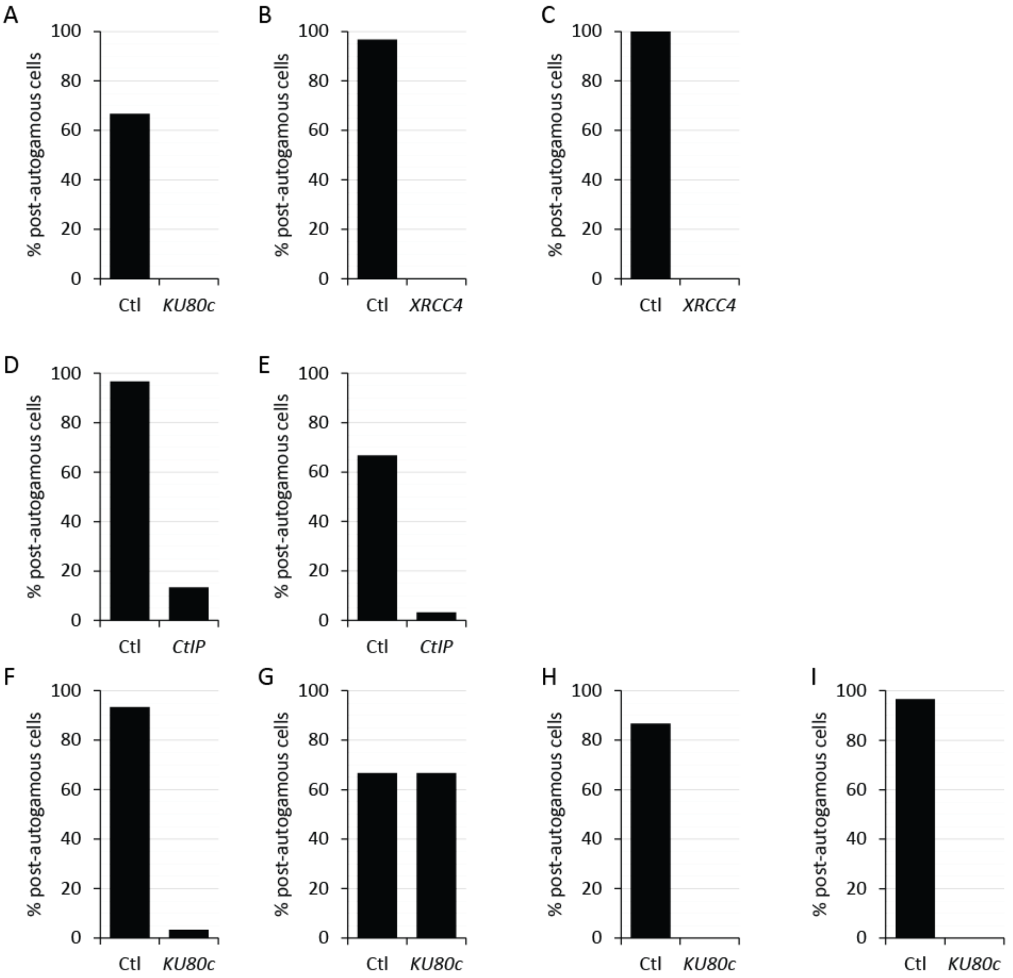


**Supplementary Figure S1. Survival of post-autogamous sexual progeny in the different RNAi experiments performed in this study**

Each panel shows the fraction of post-autogamous survivors with a functional new MAC, confirming the efficiency of all KDs (Ctl: control KD). **(A)** *KU80c* RNAi on cells injected with the *GFP* reporter gene (1976 cphg), used for western blot analysis (Figure 1A). **(B)** *XRCC4* RNAi experiment used for anlagen purification and DNA sequencing (Figure 1B). **(C)** *XRCC4* RNAi experiment used for western blot analysis of endogenous Pgm amounts (Figure 1D). **(D)** *CtIP* RNAi experiment used for western blot analysis of endogenous Pgm amounts (Figure 3B). **(E)** *CtIP* RNAi experiment on cells injected with the GFP reporter gene (416 cphg), used for western blot analysis (Figure 3C & Supplementary Figure S8). **(F)** *KU80c* RNAi on cells injected with the GFP reporter gene carrying the WT motif in its promoter (695 cphg), used for western blot analysis (Figure 5B). **(G)** *KU80c* RNAi on cells injected with the GFP reporter gene carrying the mutant motif in its promoter (2524 cphg), used for western blot analysis (Figure 5B). **(H)** *KU80c* RNAi on cells injected with the GFP reporter gene carrying the WT motif in its promoter (204 cphg), used for western blot analysis (Figure 5C). **(I)** *KU80c* RNAi on cells injected with the GFP reporter gene carrying the inverted motif in its promoter (188 cphg), used for western blot analysis (Figure 5C).

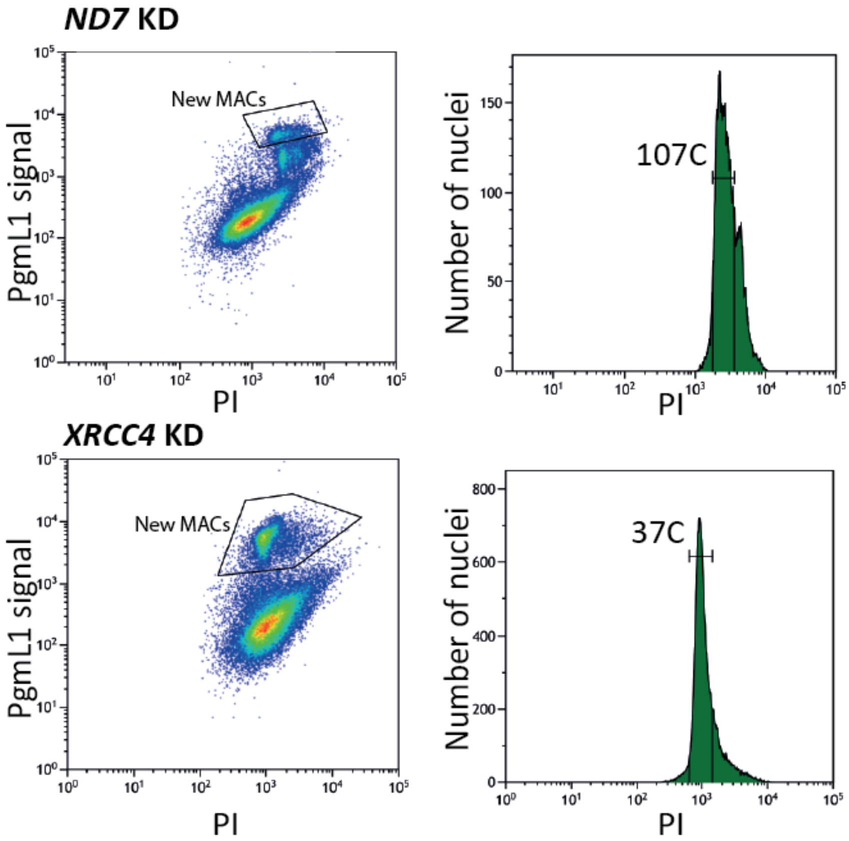


**Supplementary Figure S2. Flow cytometry on the nuclei sorted for DNA-sequencing**
(Left) Gates used to sort the new MACs from nuclei extracted at T30 of cell culture in control KD (*ND7* KD) or *XRCC4* KD. (Right) Evaluation of the ploidy of the new MACs.

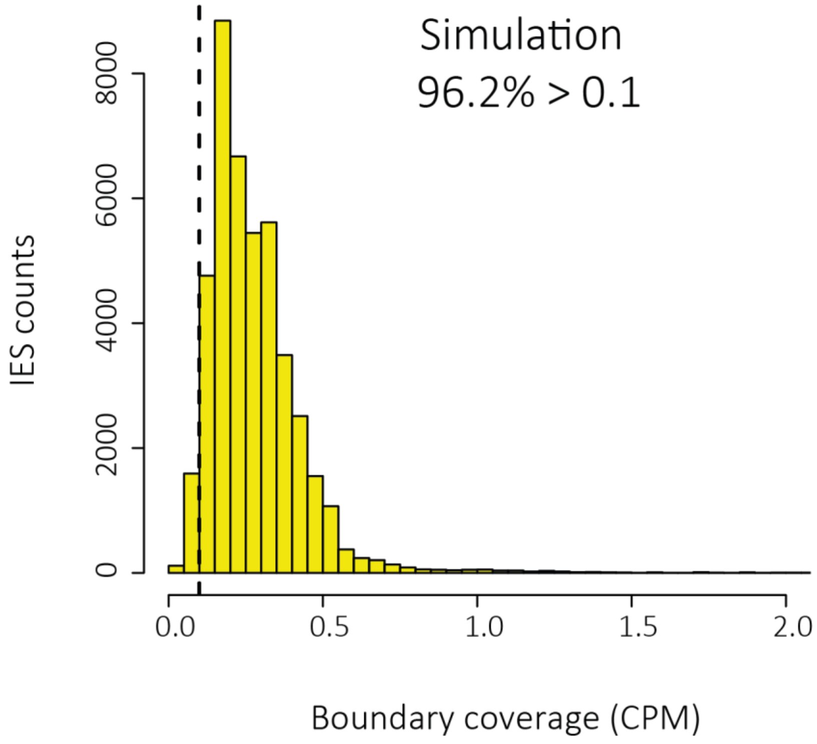


**Supplementary Figure S3. IES boundary coverage distribution for a simulated dataset with 100% IES retention.**

A simulated sequencing dataset was generated from the MAC+IES genome using ART_Illumina. The left boundary coverage was estimated for each IES using the ParTIES MIRET module (Denby Wilkes *et al*, 2015) and normalized by total simulated read coverage per Mbp (CPM). The vertical dotted line indicates the >0.1 CPM threshold.

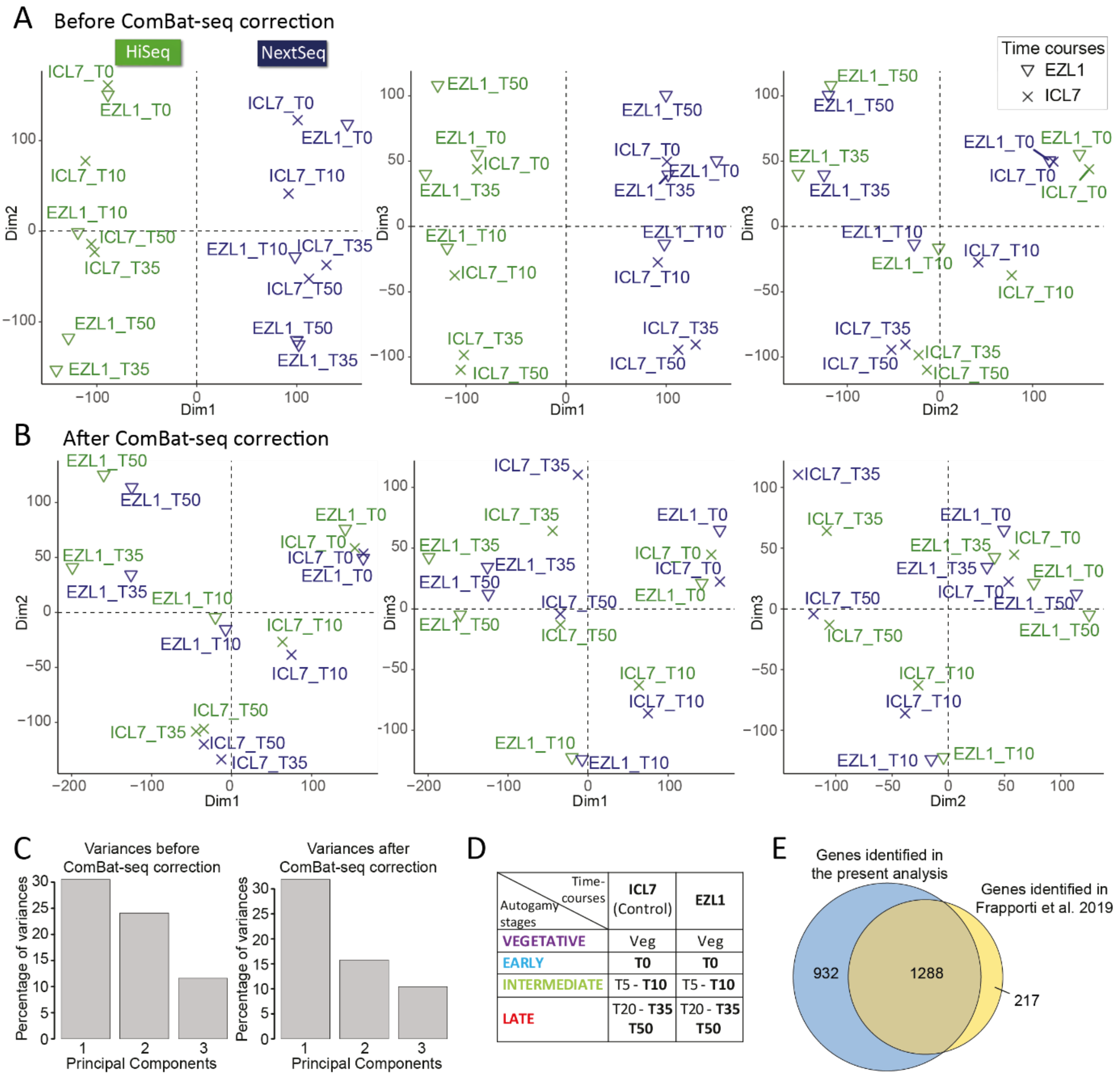


| Autogamy stages \ Time-courses | ICL7 (Control) | EZL1 |
|---|---|---|
| VEGETATIVE | Veg | Veg |
| EARLY | **T0** | **T0** |
| INTERMEDIATE | T5 - **T10** | T5 - **T10** |
| LATE | T20 - **T35** **T50** | T20 - **T35** **T50** |

**Supplementary Figure S4. Validation of ComBat-seq using *ICL7* (control) and *EZL1* RNAi samples**
**(A)** & **(B)** HiSeq (green) and NextSeq (dark blue) RNA sequencing data were obtained at T0, T10, T35 and T50 during ICL7 and EZL1 autogamy time-courses (Frapporti *et al*, 2019; Miró-Pina *et al*, 2022). We extracted the vst (variance stabilizing transformation)-normalized counts from DESeq2 analyses performed before (A) or following (B) ComBat-seq correction (Zhang *et al*, 2020) (sva 3.38.0 R package). Three dimensions of the PCA are shown. **(C)** Percentages of variances for the PCAs shown in A & B. **(D)** Table showing the 8 groups of replicates used for DE analysis, after manual curation and collapse of technical replicates (in bold). **(E)** Venn diagram showing the comparison of the set of previously identified overexpressed genes at the LATE stage (Frapporti *et al*, 2019) with those found in the present analysis, with a fold-change significance threshold >2.

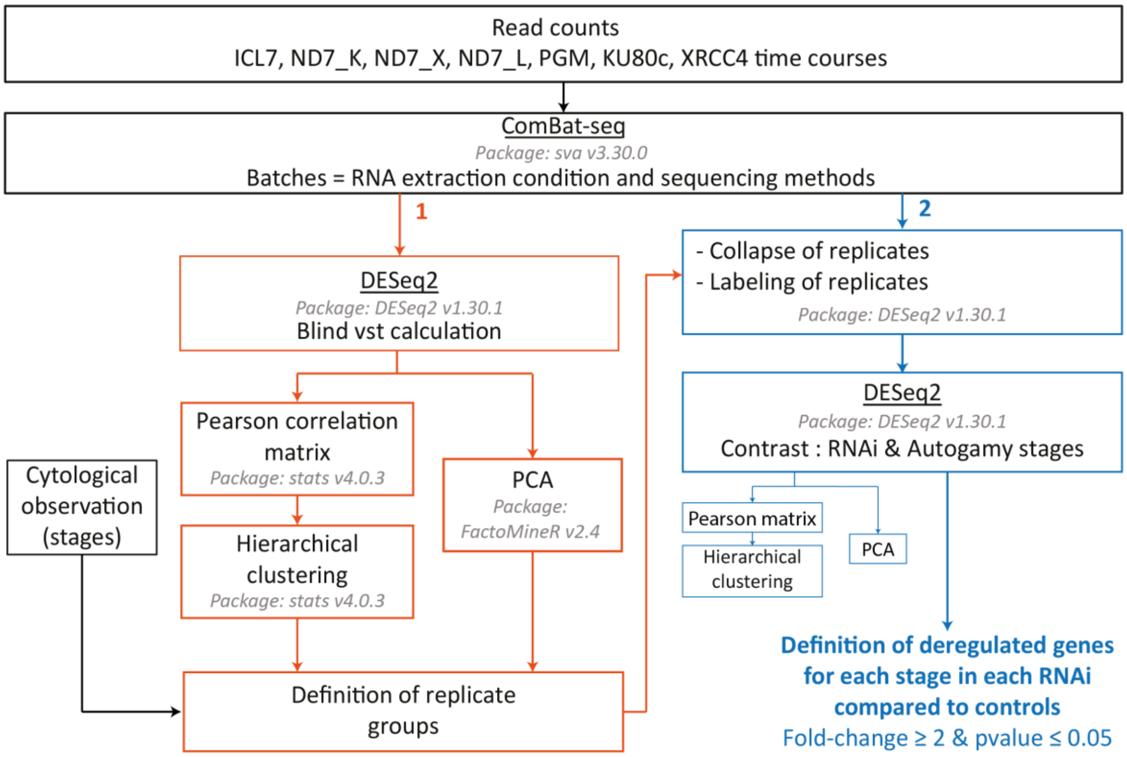


**Supplementary Figure S5. Transcriptome analysis workflow for the identification of deregulated genes**

All read counts from the indicated autogamy time-courses were first corrected for batch effects using ComBat-seq. (1) We performed a first DESeq2 analysis using the corrected data and extracted the vst-normalized values. Groups of replicates were defined using both Pearson correlation followed by hierarchical clustering, and PCA. We manually curated the groups using cytological observations of autogamy stages (Supplementary Figure S6). (2) When appropriate, we collapsed the time-points that were sequenced twice, before labelling the replicates and running a second DESeq2 round. Replicate groups were confirmed using PCA and hierarchical clustering of vst-normalized counts. The results were extracted using RNAi conditions and autogamy stages (VEGETATIVE, EARLY, INTERMEDIATE, LATE) as contrast to define deregulated genes in each RNAi relative to control conditions, at each autogamy stage.

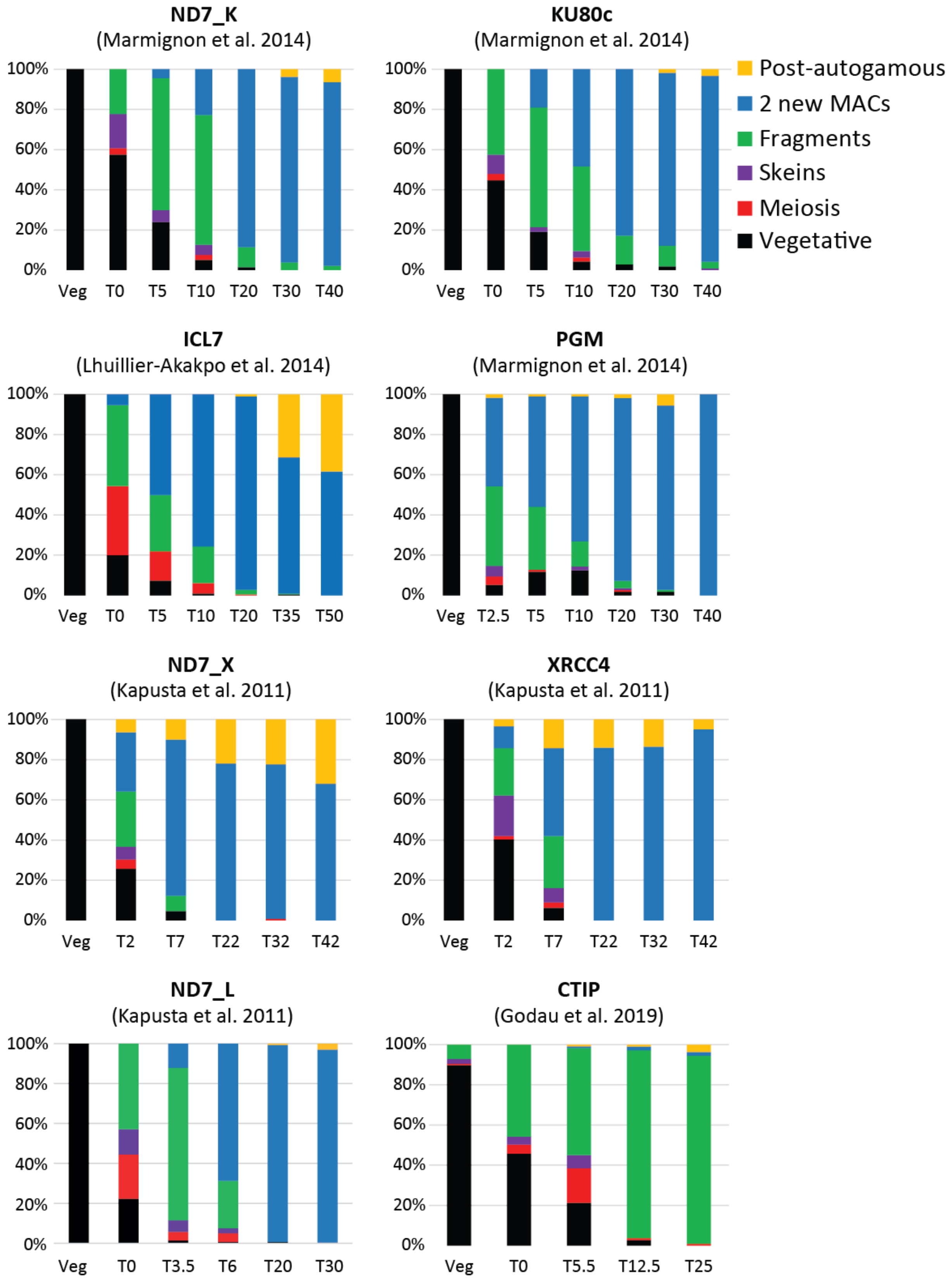


**Supplementary Figure S6. Progression of autogamy stages in the different time-courses**
All data were extracted from previously published work (Marmignon *et al*, 2014; Lhuillier-Akakpo *et al*, 2014; Kapusta *et al*, 2011; Godau *et al*, 2019). References are indicated above each diagram.

A

| | Controls | | | | | | | |
|---|---|---|---|---|---|---|---|---|
| Time courses / Autogamy stages | ICL7 | ND7_K | ND7_X | ND7_L | CTIP | XRCC4 | PGM | KU80c |
| VEGETATIVE | Veg | Veg | Veg | Veg | Veg | Veg | Veg | Veg |
| EARLY | **T0** | T0 | T2 | T0 | T0 | T2 | - | T0 |
| INTERMEDIATE | T5 - **T10** | T5 - T10 | T7 | T3.5 - T6 | T5.5 - T12.5 | T7 | T2 - T5 - T10 | T5 - T10 |
| LATE | T20 - **T35** - **T50** | T20 - T30 - T40 | T22 - T32 - T42 | T20 - T30 | T25 | T22 - T32 - T42 | T20 - T30 - T40 | T20 - T30 - T40 |

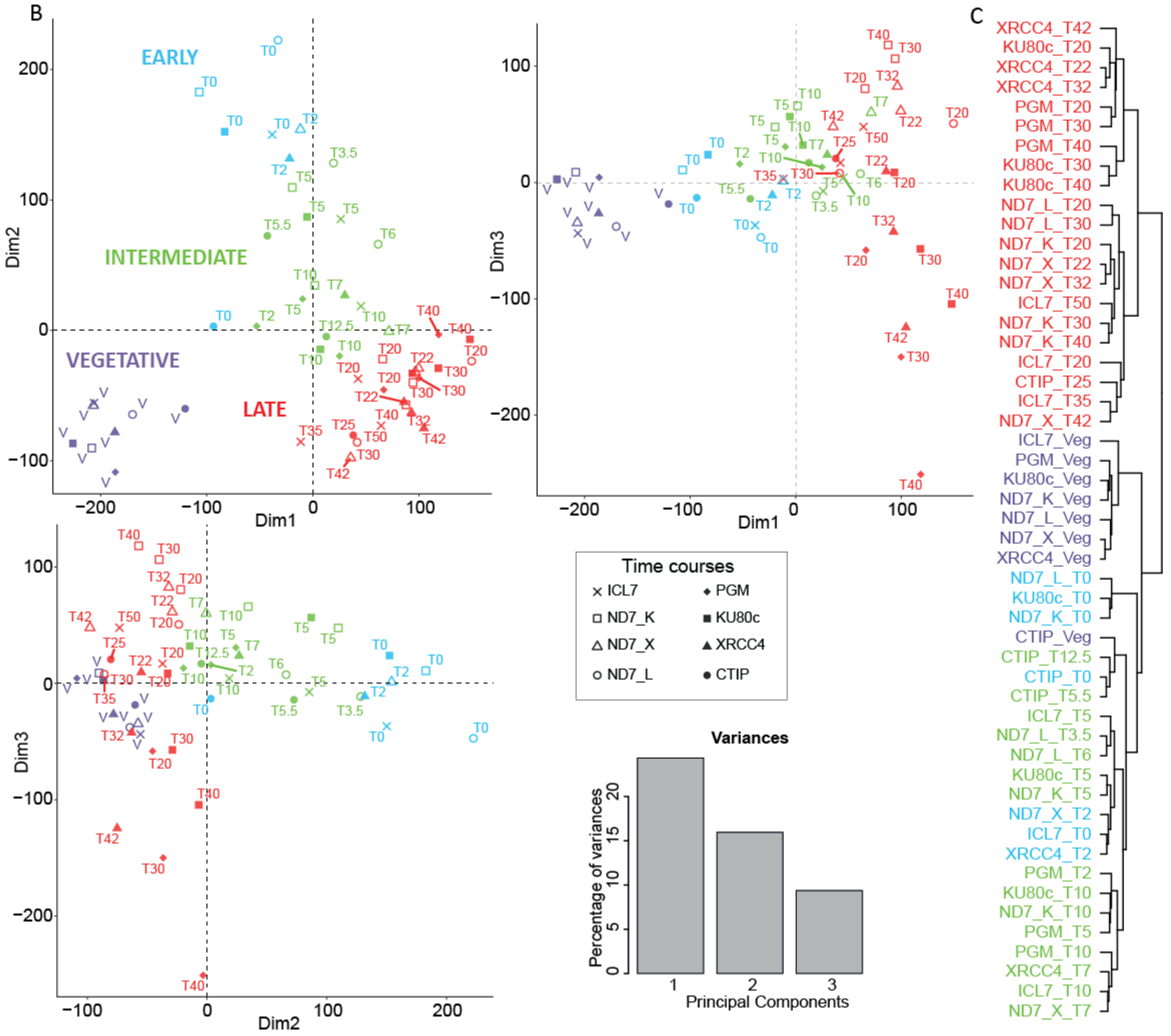


**Supplementary Figure S7. Definition of the groups of biological replicates**

**(A)** Table showing the 19 groups of replicates used for DE analysis, after manual curation and collapse of technical replicates for the ICL7 time-course (in bold). For each autogamy stage, all controls are grouped as biological replicates. **(B)** Representation of the first 3 dimensions of the PCA of the vst normalized data extracted from the second round of DESeq2 (see part 2 of Supplementary Figure S). **(C)** Hierarchical clustering (Pearson correlation matrix) of all datasets used in B (same color code). In B and C, the different time-points are colored according to the autogamy stages defined in A (VEGETATIVE, EARLY, INTERMEDIATE & LATE).

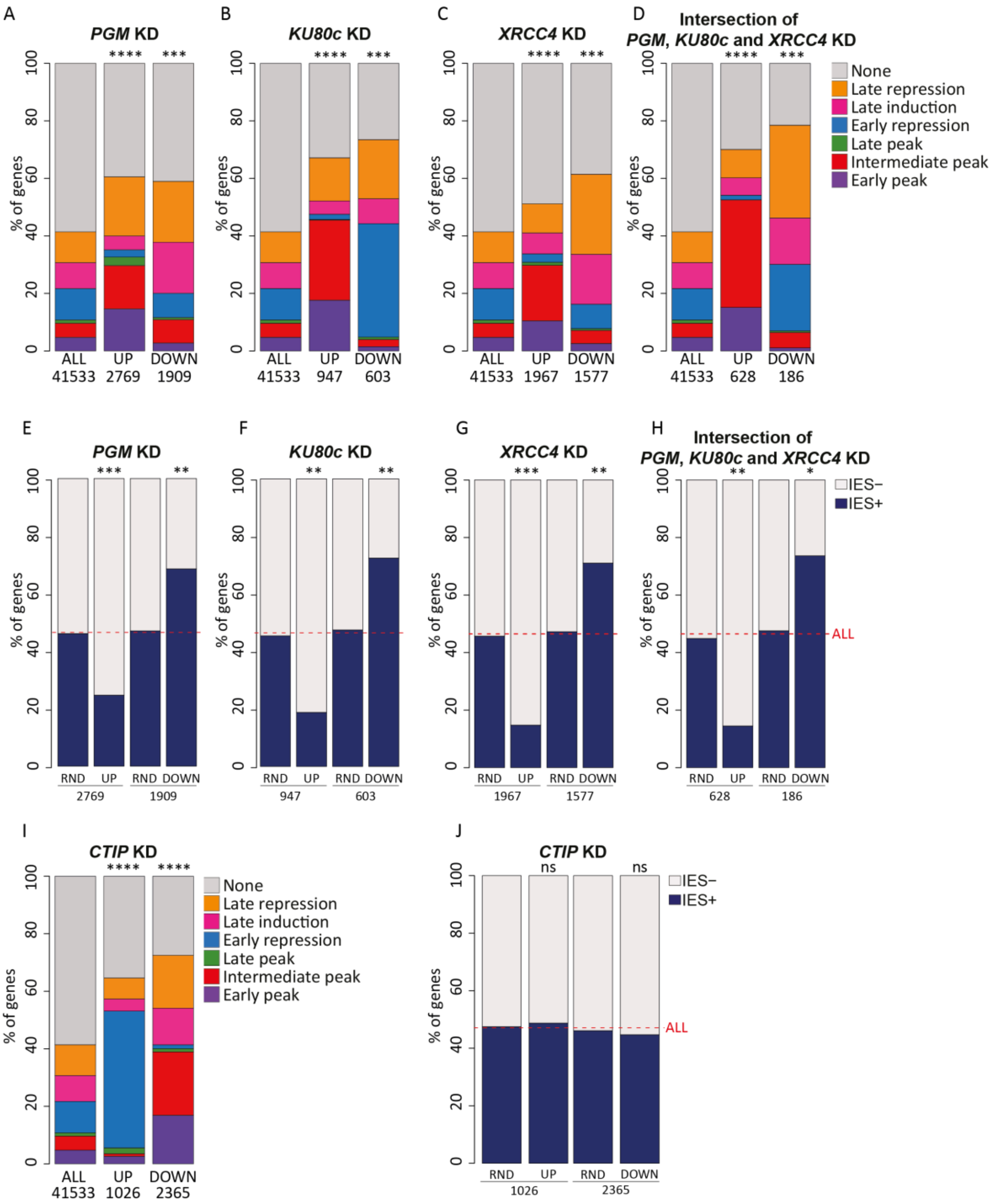


**Supplementary Figure S8. Proportions of gene expression clusters and genes with IESs in their MIC version among the combinations of deregulated genes in the different KDs**

(A-H) Proportion of each gene expression cluster (A-D) and % of genes with IES (E-H) among deregulated genes at LATE stage, in each KD and the combination of the three KDs. (I-J) Proportion of each gene expression cluster (I) and genes with IESs (J) among the deregulated genes at EARLY or INTERMEDIATE stage in *CtIP* KD. In E-H & J: the horizontal red dotted line represents the % of all genes with at least one IES (ALL), which is used as a reference for statistical analyses. For each set of UP or DOWN genes, RND shows the mean % of IES+ genes from 1000 random samples of the same number of genes, with similar proportions of each expression cluster.

****: p-value $<10^{-200}$; ***: p-value $<10^{-100}$; **: p-value $<10^{-20}$; *: p-value $<10^{-10}$; ns (not significant): p-value >0.05 (Chi$^2$ test).

A

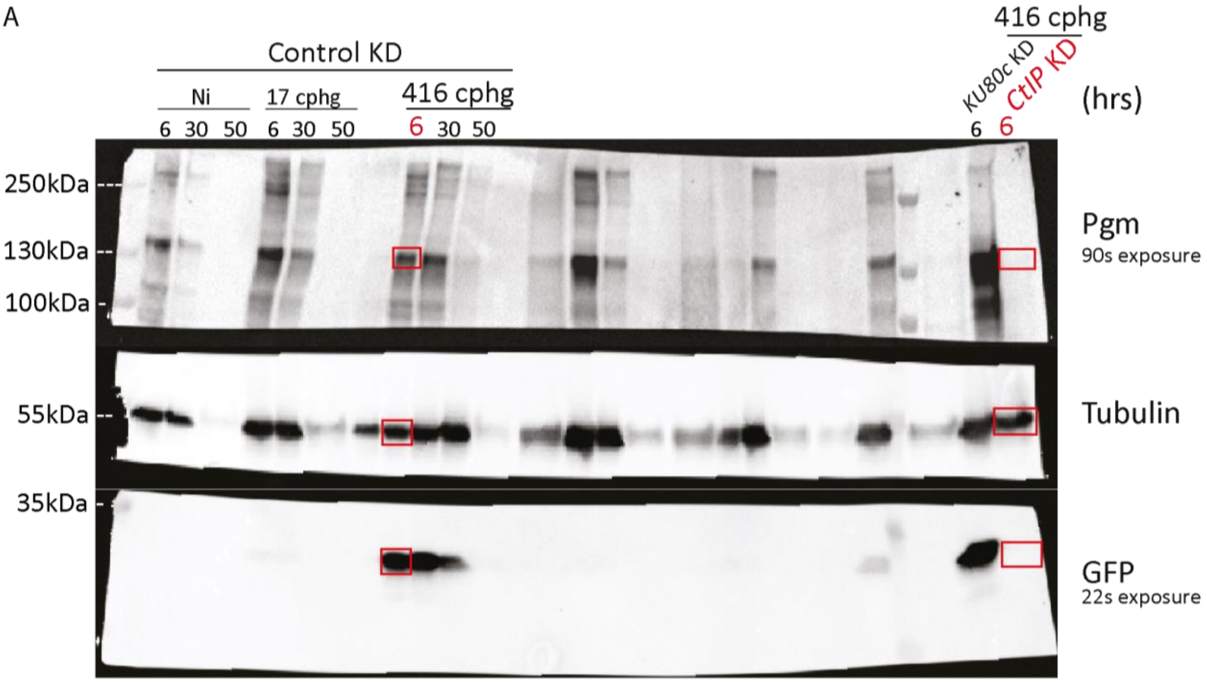


B

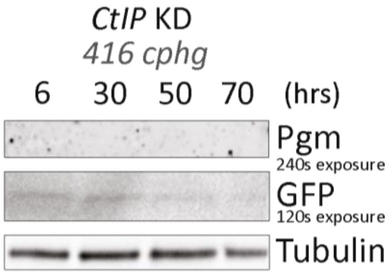


**Supplementary Figure S9. Western blot analysis of Pgm downregulation in a *CtIP* KD**

**(A**) Full-size image of the membrane used in Figure 3C. The progression of autogamy was followed for different injected clones carrying the GFP reporter construct. The samples corresponding to the clone shown in Figure 3C (416 cphg) are boxed in red for the control and *CtIP* KD. The membrane was cut in three to simultaneously monitor the production of Pgm (top), Tubulin (middle) and GFP (bottom) using appropriate antibodies. **(B)** Western blot analysis of Pgm and GFP production during autogamy in a *CtIP* RNAi for the injected clone shown in A.

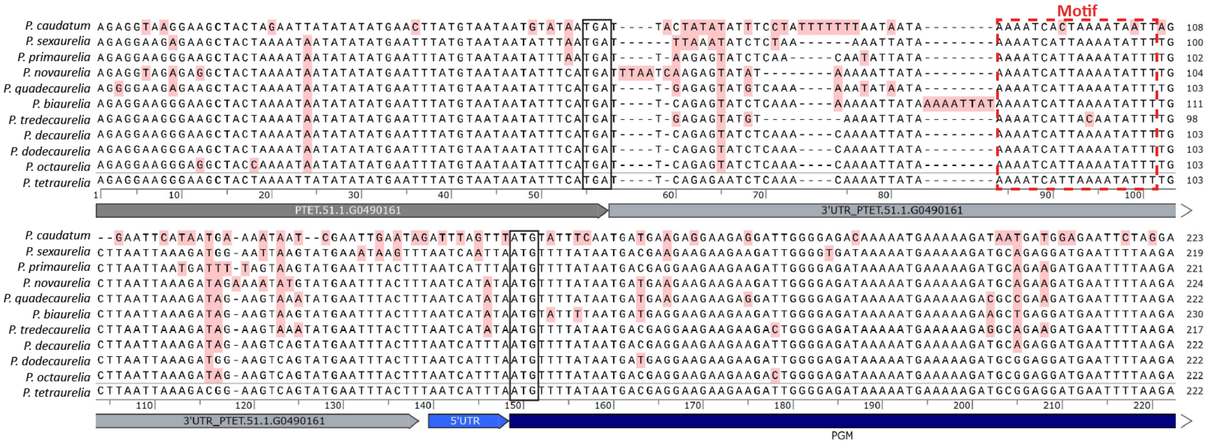

**Supplementary Figure S1. Alignment of the nucleotide sequences of the *PGM* gene start and its upstream region from 11 *Paramecium* species.**

Sequences were extracted from ParameciumDB (Arnaiz *et al*, 2019), based on (Gout *et al*, 2019; McGrath *et al*, 2014b, 2014a). The *P. tetraurelia* sequence is annotated below the alignment and the nucleotide substitutions found in other species are highlighted in pink. The conserved motif is boxed in red, the *PGM* start codon and the stop codon of the upstream gene are boxed in black.

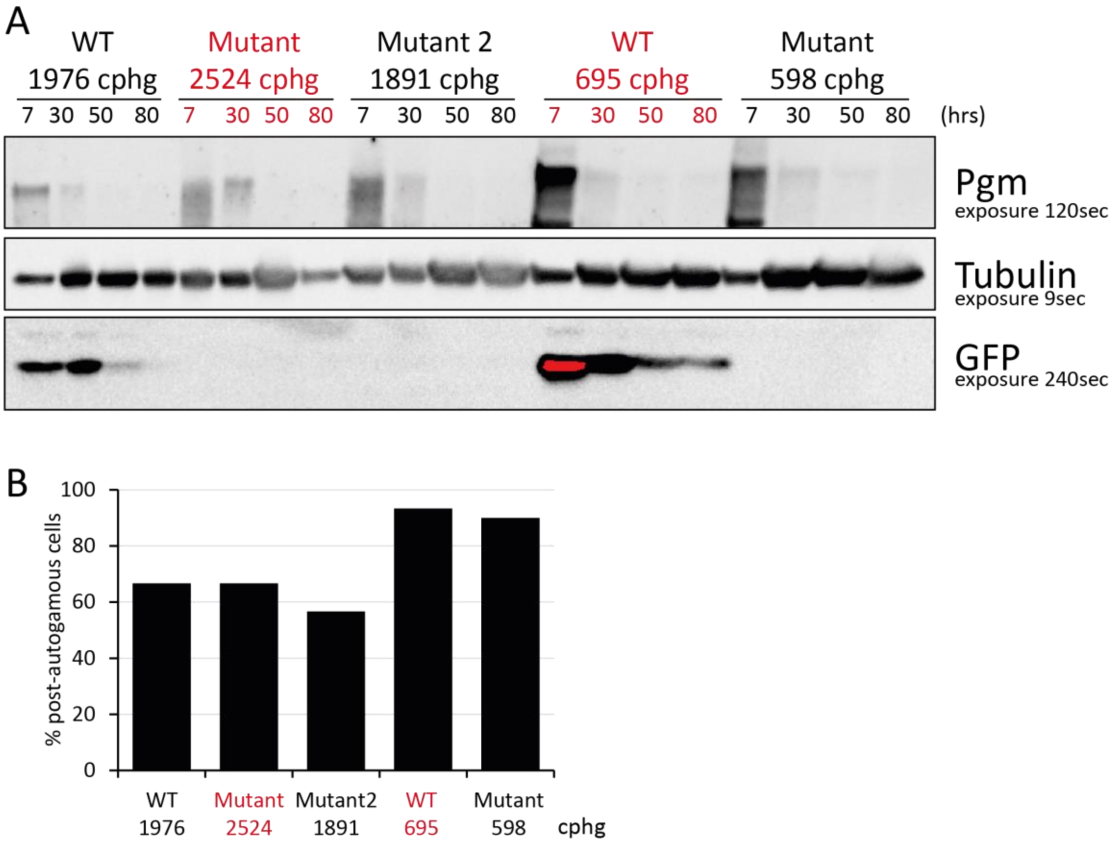


**Supplementary Figure S11. Effect of the mutated motif on the *GFP* reporter transgene expression**
**(A)** Western blot quantification of the levels of endogenous Pgm and GFP expressed from the reporter constructs injected in different clones at various cphg, during an autogamy time-course of cells subjected to control RNAi. The promoters of the *GFP* reporter transgenes carry the wild-type motif (WT: AAAATCATTAAAATATTT) or its mutated versions (mutant: AACATTAATAAAATATTT, mutant 2: AATGATTTTAAAATATTT, the mutated bases of the motif are underlined). Time-points are in hours (hrs) following T0. The time courses in red correspond to those presented in the Figure 5B. In the GFP panel (bottom), the red label indicates that the intensity of the signal was saturating. **(B)** Survival tests of each clone presented in panel A after a control RNAi.

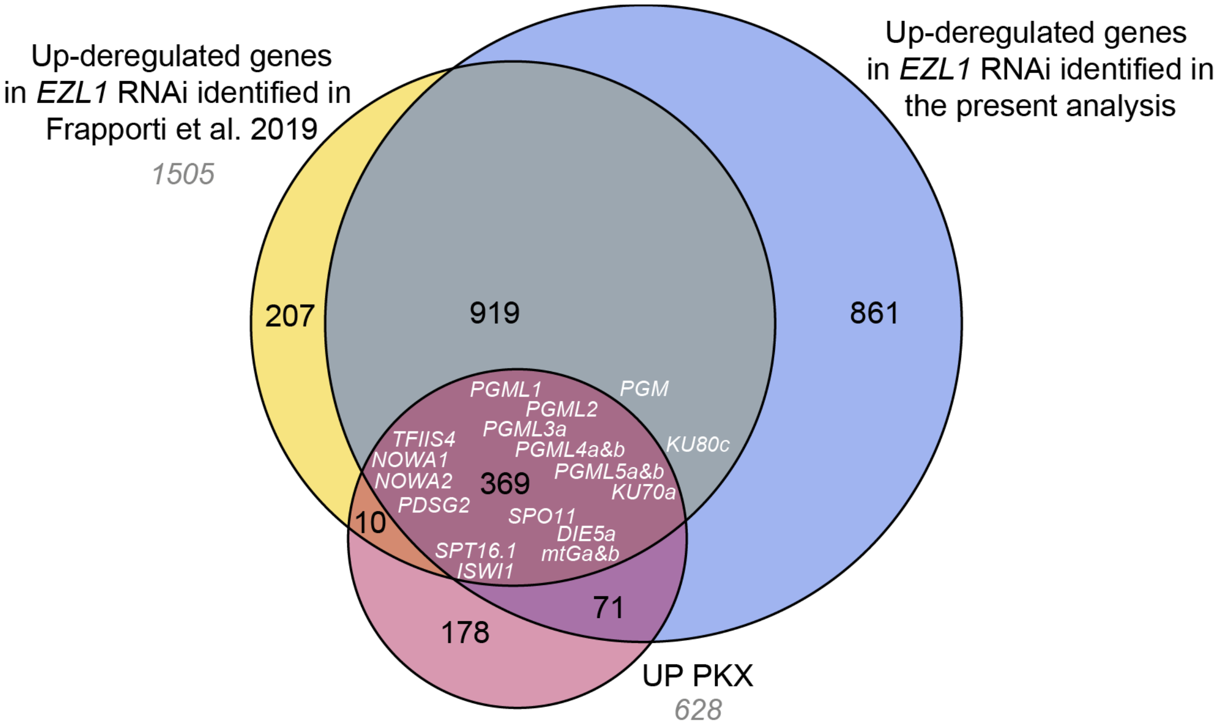


**Supplementary Figure S12. Genes overexpressed in *PGM*, *KU80c*, *XRCC4* and *EZL1* RNAi.**
Venn diagram showing the intersection of the genes identified as overexpressed in *PGM*, *KU80c* and *XRCC4* RNAi (UP PKX) with the genes overexpressed in *EZL1* RNAi according to (Frapporti *et al*, 2019) and this study. The 18 known UP PKX genes and the *PGM* and *KU80c* genes are indicated in white.

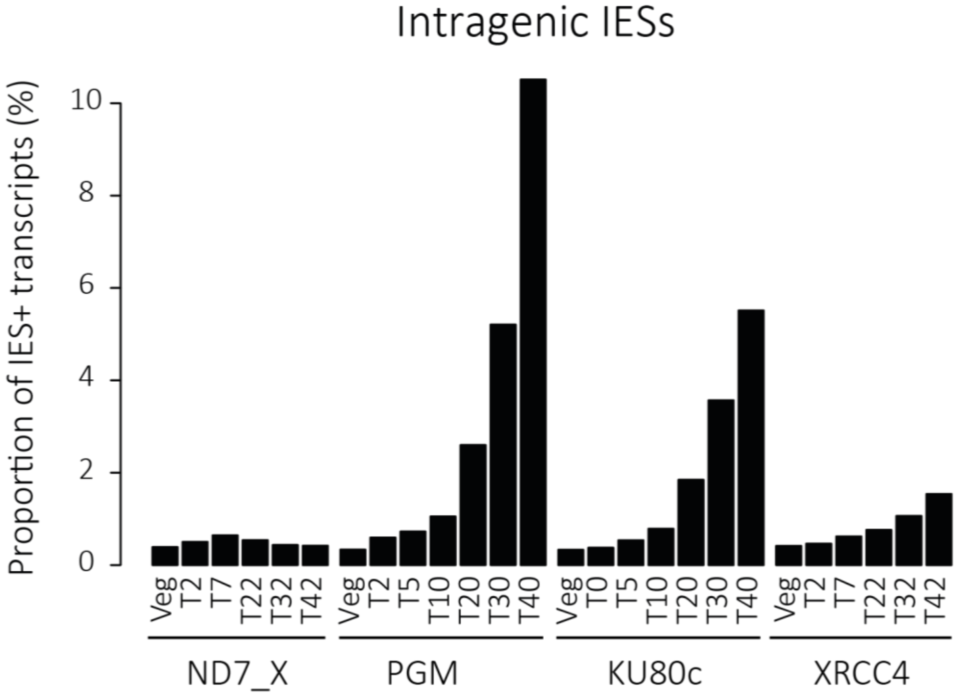


**Supplementary Figure S13. Percentage of IES+ transcripts produced by the new MACs during autogamy in control, *PGM*, *KU80c* and *XRCC4* KDs.**

For each sample, the percentage of sequencing reads covering the left boundary (IES+ reads)of the 37969 intragenic IESs was calculated relative to the sum of IES+ and IES- reads covering the same set of intragenic IESs. Relative to the control (ND7_X time-course), lower amounts of IES+ transcripts accumulate in the XRCC4 time-course than in PGM or KU80c time-courses. This may in part be explained by the persistence of unrepaired DSBs at the excision sites of a fraction of IESs, which prevents genes from being assembled in the new MACs.
